# A Non-redundant Role for T cell-derived IL-22 in Antibacterial Defense of Colonic Crypts

**DOI:** 10.1101/2021.05.11.443677

**Authors:** Carlene L. Zindl, Steven J. Witte, Vincent A. Laufer, Min Gao, Zongliang Yue, Daniel J. Silberger, Stacey N. Harbour, Jeffrey R. Singer, Duy Pham, Carson E. Moseley, Baiyi Cai, Henrietta Turner, Fran E. Lund, Bruce A. Vallance, Alexander F. Rosenberg, Jake Y. Chen, Robin T. Hatton, Casey T. Weaver

**Affiliations:** Departments of Pathology, University of Alabama at Birmingham, Birmingham, Alabama 35294, USA; Departments of Medicine, University of Alabama at Birmingham, Birmingham, Alabama 35294, USA; Departments of Genetics, University of Alabama at Birmingham, Birmingham, Alabama 35294, USA; Departments of Informatics Institute, University of Alabama at Birmingham, Birmingham, Alabama 35294, USA; Departments of Microbiology, University of Alabama at Birmingham, Birmingham, Alabama 35294, USA; Department of Pediatrics, University of British Columbia Vancouver, BC V6H 3V4

## Abstract

IL-22 is a key cytokine in immune defense against pathogens at barrier sites. In response to enteric attaching and effacing bacteria, IL-22 produced by type 3 innate lymphoid cells (ILC3s) is thought to be important early for induction of antimicrobial peptides (AMPs) that protect intestinal epithelial cells (IECs) in advance of T cell-derived IL-22 that arises later. Yet, the basis for a requirement for both innate and adaptive IL-22–producing immune cells in protecting the intestinal mucosa is unknown. Here, using novel mice that both report IL-22 expression and can be targeted for its lineage-specific deletion, we show that mice with deficiency of IL-22 targeted to innate immune cells, including ILC3s, have impaired STAT3 activation of surface colonic IECs colonized by bacteria early in infection. In contrast, mice with IL-22 deficiency limited to T cells have complete loss of STAT3 activation in IECs lining colonic crypts and fail to protect the crypts from bacterial invasion late despite ongoing production of IL-22 from ILC3s. T cell-derived IL-22 is required for upregulation of many host-protective genes by crypt IECs, including those encoding AMPs, neutrophil-recruiting chemokines, and mucins and mucin-related molecules, while also restricting pro-inflammatory genes downstream of IFN*γ* and TNF signals. Thus, T cell-derived IL-22 is indispensable for antibacterial defense and damage control of intestinal crypts.

## Introduction

Host defense against extracellular bacteria is orchestrated by type 3 immunity, which employs cells of the innate and adaptive immune systems that share responsiveness to IL-23 and production of IL-17 family cytokines and the IL-10 family cytokine, IL-22 (Mangan et al., 2006; Sonnenberg et al., 2011b; Zheng et al., 2008). *Citrobacter rodentium* (*C.r*) is an attaching and effacing (A/E) enteric pathogen that closely models human disease caused by enteropathogenic and enterohemorrhagic *E. coli* (EPEC and EHEC) (Collins et al., 2014; Mundy et al., 2005; Silberger et al., 2017). These Gram-negative bacteria use a type III secretion system to inject effector molecules via the apical surface of intestinal epithelial cells (IECs), allowing them to attach and efface the microvilli of host IECs and establish colonization (Donnenberg et al., 1997; Frankel et al., 1998). Clearance of *C.r* from the epithelial surface occurs when bacterial-laden IECs are shed into the lumen (Barker et al., 2008; Clevers, 2013). However, A/E pathogens have evolved mechanisms to inhibit apoptosis and turnover of IECs to which they adhere (Hemrajani et al., 2010; M. Kim et al., 2010; Nougayrède et al., 2005), thereby prolonging colonization. Moreover, *C.r* manipulates host IEC metabolism to create an environment suitable for growth and evasion from innate immune responses (C. N. Berger et al., 2017). Therefore, antigen-specific CD4 T-cell and B-cell responses are ultimately required for pathogen eradication (Bry et al., 2005; Maaser et al., 2004; Simmons et al., 2003; Vallance et al., 2002).

A histopathological hallmark of *C.r* infection is elongation of the crypt epithelium (hyperplasia) and goblet cell depletion (hypoplasia) in the large intestine, or colon (C. N. Berger et al., 2017; Bergstrom et al., 2008; Borenshtein et al., 2009; Chan et al., 2013; Ma et al., 2006; Papapietro et al., 2013), the distal part of which is the infectious niche of *C.r* in mice as opposed to the ileum and transverse colon of *E. coli* (EPEC and EHEC respectively) in humans (Croxen et al., 2013). Elongation of the colonic crypts is thought to isolate intestinal stem cells (ISCs) that reside in the base of crypts from physical and metabolic damage that results from infection, thereby preventing destruction of the progenitors that give rise to all cells of the intestinal epithelial monolayer (Kaiko et al., 2016; Y. Liang et al., 2017; Matsuki et al., 2013; Okada et al., 2013). Accelerated production of IEC progenitors, or transient-amplifying (TA) cells correlates with increased shedding of *C.r*-laden luminal surface IECs (Collins et al., 2014; Higgins et al., 1999). However, details regarding mechanisms by which IEC differentiation is altered during *C.r* infection are incompletely defined.

STAT3 activation is a major output of the liganded IL-22 receptor, which is composed of IL-22Ra1 and IL-10Rb subunits and is expressed by IECs (Lindemans et al., 2015). IL-22/IL-22R/STAT3 signaling into IECs has been shown to be important for mucosal barrier protection and restitution of the intestinal epithelium during infection (Basu et al., 2012; Pickert et al., 2009; Wittkopf et al., 2015; Zheng et al., 2008). IL-22R signaling upregulates various host defense molecules, such as antimicrobial peptides (e.g., Reg3 and S100a family members) (S. C. Liang et al., 2006; Wolk et al., 2006; Zheng et al., 2008), proteins implicated in inflammatory responses (e.g., Lbp, Saa, complement and chemokines) (Aujla et al., 2008; Boniface et al., 2005; Hasegawa et al., 2014; S. C. Liang et al., 2010), and proteins involved in altering the mucus layer (e.g., Muc1 and Fut2) (Pham et al., 2014; Sugimoto et al., 2008). Several different types of innate immune cells, including ILC3s, NK cells, NKT cells, DDT cells and neutrophils (Cella et al., 2008; 2010; Chen et al., 2016; Colonna, 2009; J. S. Lee et al., 2011; Y.-S. Lee et al., 2015; Satpathy et al., 2013; Sonnenberg et al., 2011b; 2011a; Spits et al., 2013; Zheng et al., 2008; Zindl et al., 2013) can respond to IL-23 from dendritic cells to produce IL-22 that acts on IECs. CD4 T cells of the Th17 pathway—Th17 and Th22—also produce IL-22, whether induced in response to IL-23 or TCR signaling (Akdis et al., 2012; Basu et al., 2012; Guo, 2016; C. J. Kim et al., 2012; Trifari et al., 2009). ILC3s are thought to be the major innate cell population expressing IL-22 during *C.r* infection and are crucial for early host protection (Rankin et al., 2015; Sonnenberg et al., 2011b; Spits et al., 2013). Th17 and Th22 cells induced during *C.r* infection contribute to IL-22 following recruitment to the intestinal mucosa later in *C.r* infection (S. C. Liang et al., 2006; Zheng et al., 2006). These cells appear to be important in enhancing barrier protection and limiting epithelial cell damage as infection progresses (Basu et al., 2012; Silberger et al., 2017; Zenewicz et al., 2008). In addition, recent studies show that T cell-dependent, *C.r*-specific IgG is required to eradicate virulent *C.r* (Kamada et al., 2015). However, the relative contributions of innate immune cells and CD4 T cells and the mechanisms by which IL-22-producing T cells control *C.r* infection are unclear.

Using new IL-22 reporter/conditional knockout mice with which to identify IL-22 producers and target deficiency of IL-22 to different cell populations, we find that ILC3s and Th17/Th22 cells have distinct roles in activating IECs during *C.r* infection. Whereas innate cell-derived IL-22 dominates first and targets superficial IECs early in infection to limit the initial wave of *C.r* colonization and spread, T cell-derived IL-22 is indispensable later for induction of heightened and sustained STAT3 activation in both superficial and crypt IECs to prevent bacterial invasion of colonic crypts and limit bacterial dissemination as infection progresses. RNA-seq analysis of colonic IECs indicate that IL-22-producing T cells uniquely mobilize multiple mechanisms that underlie their essential role in protecting the crypts and preserving ISCs that provide progeny for restitution of the infected intestinal epithelium.

## Results

### Distinct spatiotemporal distribution of IL-22–producing innate and adaptive immune cells during *C.r* infection

It is clear from our own and others’ studies that multiple immune cell types can produce IL-22 in the intestines (Basu et al., 2012; Cella et al., 2008; Colonna, 2009; Silberger et al., 2017; Sonnenberg et al., 2011b; Spits et al., 2013; Trifari et al., 2009; Zindl et al., 2013). To better characterize dynamics of the location and number of IL-22–producing cells in the intestines during the course of *C.r* infection, we developed gene-targeted IL-22 reporter/conditional knockout (cKO) mice to track and/or delete specific subsets of IL-22–producing immune cells (*Il22*^hCD4.fl^ mice, hereafter referred to as *Il22*^hCD4^; **Figures S1A-S1D**). Using the reporter function of these mice, we found that at steady state in the colons of naïve mice type 3 innate lymphoid cells (ILC3s) were the dominant IL-22^+^ cells (**Figures 1A-1E**). During early *C.r* infection (d3-6), mCD4^+^TCRβ^−^ ILC3s (LTi cells) produced the greatest amount of IL-22 in the colon (**Figures 1A-1C**) and cecum (data not shown) on a per-cell basis, although comparable numbers of IL-22^+^ mCD4^−^TCRβ^−^ cells (non-LTi ILC3s) were present. Interestingly, no significant change in the numbers of IL-22^+^ ILCs was observed during the course of the infection, suggesting that either these cells do not proliferate during infection or turnover so as to be replaced at the same rate they are lost during infection. This is consistent with recent reports that ILCs populate non-lymphoid tissues early in life and remain largely static in these sites (Ahlfors et al., 2014; Gasteiger et al., 2015) (**Figures 1A** and **1B**). In contrast, after the first week post-inoculation, the numbers of IL-22^+^ CD4 T cells in the infected mucosa increased rapidly and ultimately outnumbered all of the IL-22^+^ innate cells combined (**Figures 1A** and **1B**). Thus, the total number of colonic IL-22^+^ ILCs is largely static throughout the course of infection, whereas the ingress of IL-22^+^ CD4 T cells rapidly increase in the colon during the later phase of infection (from day 6 on), whether due to migration into the colon, local proliferation or both.

**Figure 1.**
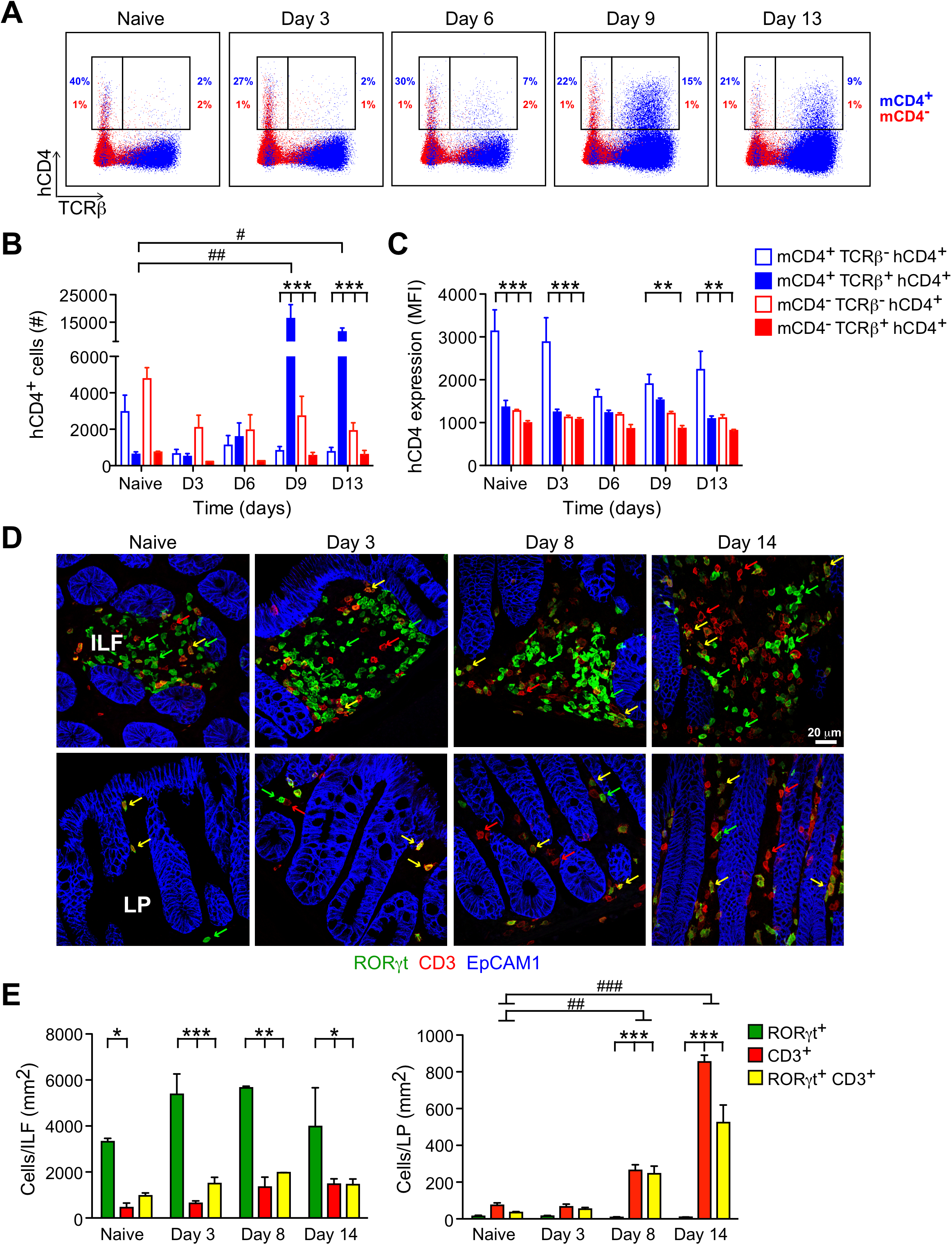
Dynamics of IL-22 expression and cellular localization during *C.r* infection. (A) Colon LP cells were isolated from naïve and *C.r*-infected *Il22*^hCD4^ mice, stimulated with PMA/Ion + rmIL-23 for 4 hrs and then stained for TCRβ, hCD4, mCD4, and L/D dye and analyzed by flow cytometry. Numbers represent percentages of hCD4 (IL-22^+^) innate cells (TCRβ^−^) or T cells (TCRβ^+^) and are fractionated into CD4^+^ (blue) and CD4^−^ (red). (B) Cell numbers and (C) IL-22/hCD4 expression based on MFI. Populations were fractionated into mCD4^+^ (open; blue) and mCD4^−^ (open; red) innate cells, and mCD4^+^ (solid; blue) and mCD4^−^ (solid; red) T cells. (D) Colons from naïve and *C.r*-infected *Rorc*^EGFP^ mice were stained for GFP (green), CD3 (red), and EpCAM1 (blue). Green arrows depict RORD/GFP^+^ cells, red arrows depict CD3^+^ cells, and yellow arrows depict GFP^+^ CD3^+^ cells. (E) Quantitation of cells in colonic ILFs and LP. One-way ANOVA; ^#^*p* <0.05, ^##^*p* <0.01 and ^###^*p* <0.001 comparing naïve and infected mice. Two-way ANOVA; \**p* <0.05, \*\**p* <0.01 and \*\*\**p* <0.001 comparing different cell populations. 3-4 mice per time point, 3 independent experiments.

Because *in situ* detection of hCD4/IL-22 expression by immunostaining proved unreliable using available antibodies, a *Rorc*/EGFP BAC reporter mouse line (Lochner et al., 2008) was used to identify and localize colonic ROR*γ*t^+^ cells, including IL-22^+^ ILC3s and Th17/Th22 cells (**Figures 1D** and **1E** and **Figures S1E-S1G**). Notably, while some IL-22^+^ROR*γ*t^+^ cells were found within the intestinal epithelium during infection, the great majority of IL-22^+^ROR*γ*t^+^ cells were found within non-epithelial tissue compartments, and the ILC3s present were largely NKp46^−^ (**Figures S1E-S1G**). Moreover, in naïve mice and during *C.r* infection, ROR*γ*t^+^ CD3^−^ ILC3s were largely clustered within colonic lymphoid tissues (i.e., solitary intestinal lymphoid tissues, or SILTS, and the colonic patches); very few ILC3s were found in the lamina propria (LP) and redistribution of these cells outside of ILFs did not change over the course of infection, in agreement with recent studies (Ahlfors et al., 2014; Colonna, 2018; Gasteiger et al., 2015) (**Figures 1D** and **1E**). Similarly, there were no significant changes in the small number of ROR*γ*t^+^ T cells found in ILFs in naïve mice and during infection. In marked contrast, ROR*γ*t^+^ CD4 T cells increased dramatically (>50-fold) in the LP with numerous T cells found in close apposition to intestinal crypt epithelial cells. Collectively, these findings indicate IL-22^+^ ILC3s and CD4 T cells occupy distinct microanatomic niches over the course of *C.r* infection, such that IL-22^+^ ILC3s are restricted to ILFs and static in number, whereas IL-22^+^ CD4 T cells populate the lamina propria in increasing numbers so as to become the dominant IL-22 producers. This results in more uniform distribution of IL-22–producing T cells relative to ILC3s and, in general, positions the T cells in closer proximity to the epithelial monolayer they are charged with protecting.

### IL-22 produced by either ILCs or T cells is required to restrain bacterial burden at different times during *C.r* infection

The distinct spatial and temporal deployment of IL-22^+^ ILCs and T cells during *C.r* infection suggested the possibility of complementary or unique functions for these cells in mucosal barrier defense. To evaluate their relative contributions, we crossed the *Il22*^hCD4^ cKO reporter mice with different Cre recombinase-expressing lines to target IL-22 deficiency to all immune cells, innate immune cells or T cells (**Figure 2A**). Using a bioluminescent strain of *C.r* (ICC180) that allows real-time visualization of intestinal colonization in the whole animal (Wiles et al., 2006), we found that infected global KO (gKO) mice (*EIIa*-*cre* x *Il22*^hCD4^; *Il22*^EIIa^) had a significantly increased bacterial burden compared to controls as early as 3 days after inoculation and that all gKO mice succumbed to the infection (**Figures 2B-2D**), in accord with previous results (Zheng et al., 2008). Similar to mice with global IL-22 deficiency, mice with deficiency of IL-22 targeted to innate immune cells (*Zbtb16/Plzf)-cre* x *Il22*^hCD4^; *Il22*^Plzf^ cKO)—in which *Il22* is deleted in nearly all ILCs, regardless of subset, as well as iNKT cells, *γδ* T cells (**Figure S2D**) (Constantinides et al., 2016; Kovalovsky et al., 2008; Lu et al., 2015; Savage et al., 2008)—succumbed to *C.r* infection with similar kinetics to gKO mice, correlating with a significantly heightened bacterial burden around d3 compared to *C.r*-infected control mice **(Figures 2B**, **2E** and **2F)**. However, ∼40% of *Il22*^Plzf^ cKO mice survived infection, presumably rescued by the late influx of IL-22–producing T cells not present in gKO mice, and in these mice clearance of the bacterial infection progressed with the same kinetics as WT controls over the late course of infection (**Figure 2B** and **Figures 1A** and **1B**). This was in contrast to mice with deficiency of IL-22 targeted to T cells (*Cd4-cre* x *Il22*^hCD4^; *Il22*^ΔTcell^ cKO), in which there was no increase in bacterial burden over the early course of infection, but ∼40% of mice succumbed late in association with significantly delayed clearance of bacteria compared to controls (**Figures 2B**, **2G** and **2H**). In view of our findings that *C.r* colonization of colonic IECs is detectable around d3 and crests by d5-7 after inoculation in mice (**Figures S2A-S2C**), these data establish that innate immune cell-derived IL-22 acts to limit bacterial colonization during the early phase of infection, but in the absence of an influx of IL-22^+^ T cells into the colon during the late stage of infection, is unable to further restrain the bacterial burden and prevent death of the host; thus, IL-22 derived from innate immune cells is critical to restrain *C.r* overgrowth during the early phase of infection, but is unable to compensate for T cell-derived IL-22 in bacterial restraint and host protection late.

**Figure 2.**
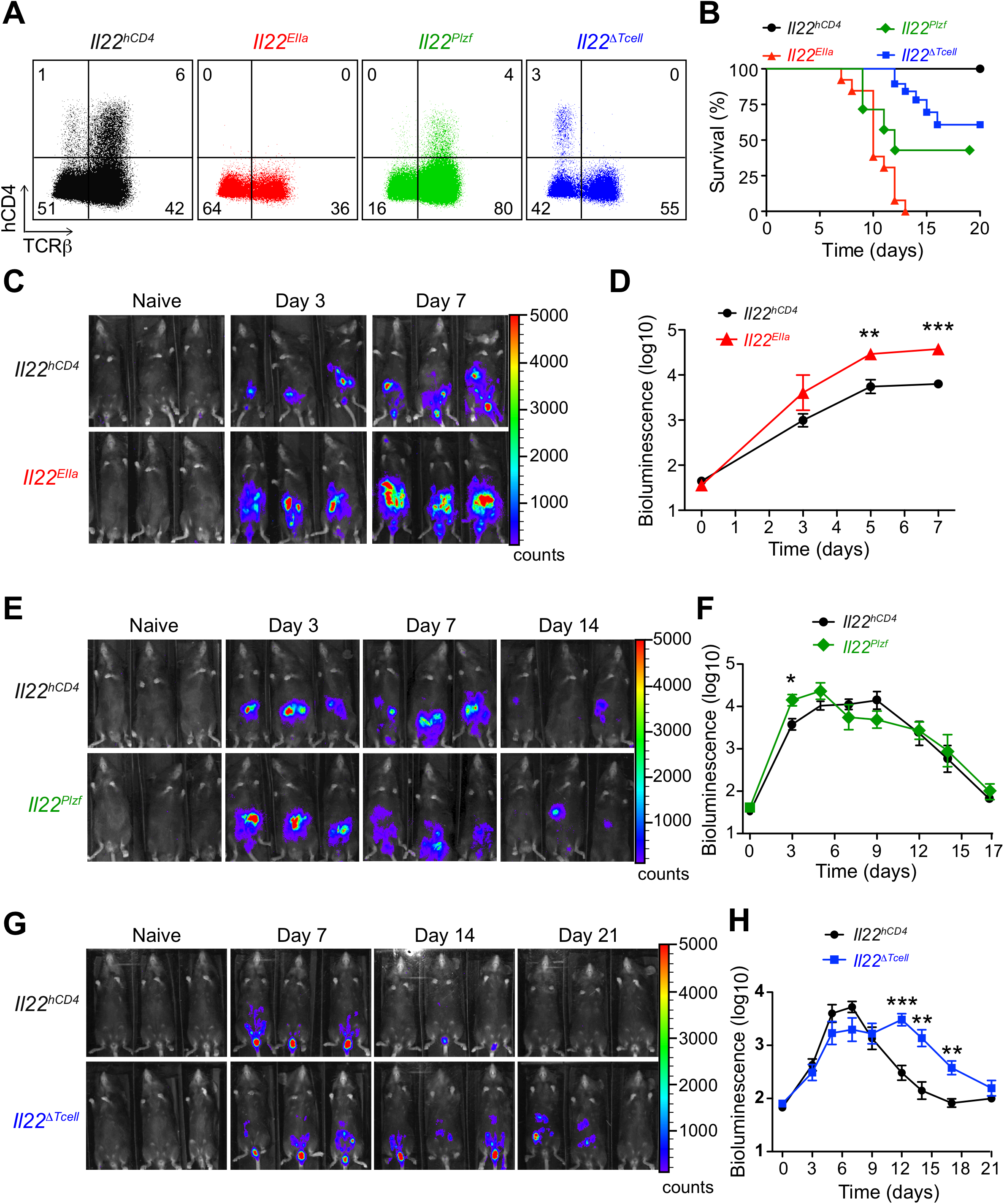
Temporal bacterial burden and fatality in *C.r*-infected *Il22* cKO mice. (A) Colon LP cells were isolated from d7 *C.r*-infected *Il22*^hCD4^ (Cntrl; black), *Il22*^EIIa^ (gKO; red), *Il22*^Plzf^ (Innate cell cKO; green), and *Il22*^ΔTcell^ (T cell cKO; blue) mice and stained for mCD4, hCD4 (IL-22), L/D dye and TCRD after stimulation with IL-23, PMA, and Ion. (B) Survival kinetics of *C.r*-infected WT, KO, and cKO mice. (C, E, G) Serial whole-body imaging and (D, F, H) Colonization kinetics of *C.r*-infected WT, KO, and cKO mice was performed on indicated days. Mann Whitney; \**p* <0.05, \*\**p* <0.01, \*\*\**p* <0.001. 4-5 mice per time point, 2 independent experiments.

### T cell-derived IL-22 is essential for protection of colonic crypts against bacterial invasion

In view of the foregoing results, we postulated that IL-22 delivered by T cells played a non-redundant role in barrier defense against *C.r* infection. To elucidate potential mechanisms by which this might occur, we examined the dynamics and distribution of bacterial epithelial attachment and the tissue response over the course of infection in each of the IL-22-deficient mouse variants. GFP-expressing *C.r* (Bergstrom et al., 2010) were used to map the density and tissue distribution of bacterial attachment to IECs and the histopathologic features of tissue injury contingent on deficiencies of IL-22 targeted to ILCs, T cell or both. Coincident with the onset of death in *C.r*-infected *Il22*^EIIa^ gKO mice (d8), we observed heightened epithelial cell damage in the cecum, middle and distal colon with significant increase in goblet cell and crypt cell loss, and depletion of crypts (**Figure 3A** and **Figure S3A**). This was accompanied by multifocal ulcerations of the mucosa and mass translocations of *C.r* bacteria (**Figure 3A**, and data not shown), consistent with our previous findings in IL-23- and IL-22-deficient mice (Basu et al., 2012; Mangan et al., 2006). During the early phase of *C.r* infection (d3/4) in control *Il22*^hCD4^ mice, focal collections of *C.r* were found attached to the luminal surface of colonic IECs (**Figure 3B**). At the same time points in *C.r*-infected *Il22*^EIIa^ gKO mice, there was uniform *C.r* colonization of the luminal surface of IECs (**Figures 3B** and **3C**), indicating that early in colonization of the epithelium, IL-22 produced by ILC3s acts to limit *C.r* colonization at the luminal surface. Notably, as infection progressed, there was more uniform distribution of *C.r* attached to the surface epithelium in control mice (d9), with extension focally into the mouth or luminal opening of the crypts, although there was no penetration deeper into the crypts (**Figure 3B**). In marked contrast, *C.r* invaded deep into colonic crypts in gKO mice by d9, coinciding with the influx of CD4 T cells into the colonic LP (**Figure 1**). Accordingly, the number of *C.r* cells attached to IECs was >10-fold higher in gKO mice compared to controls (**Figure 3C**) indicating that in the global absence of IL-22, there was loss of protection of crypts against bacterial invasion. Thus, IL-22 is required to control the progressive spread of *C.r* from small initial foci attached to surface IECs to the depths of crypts.

**Figure 3.**
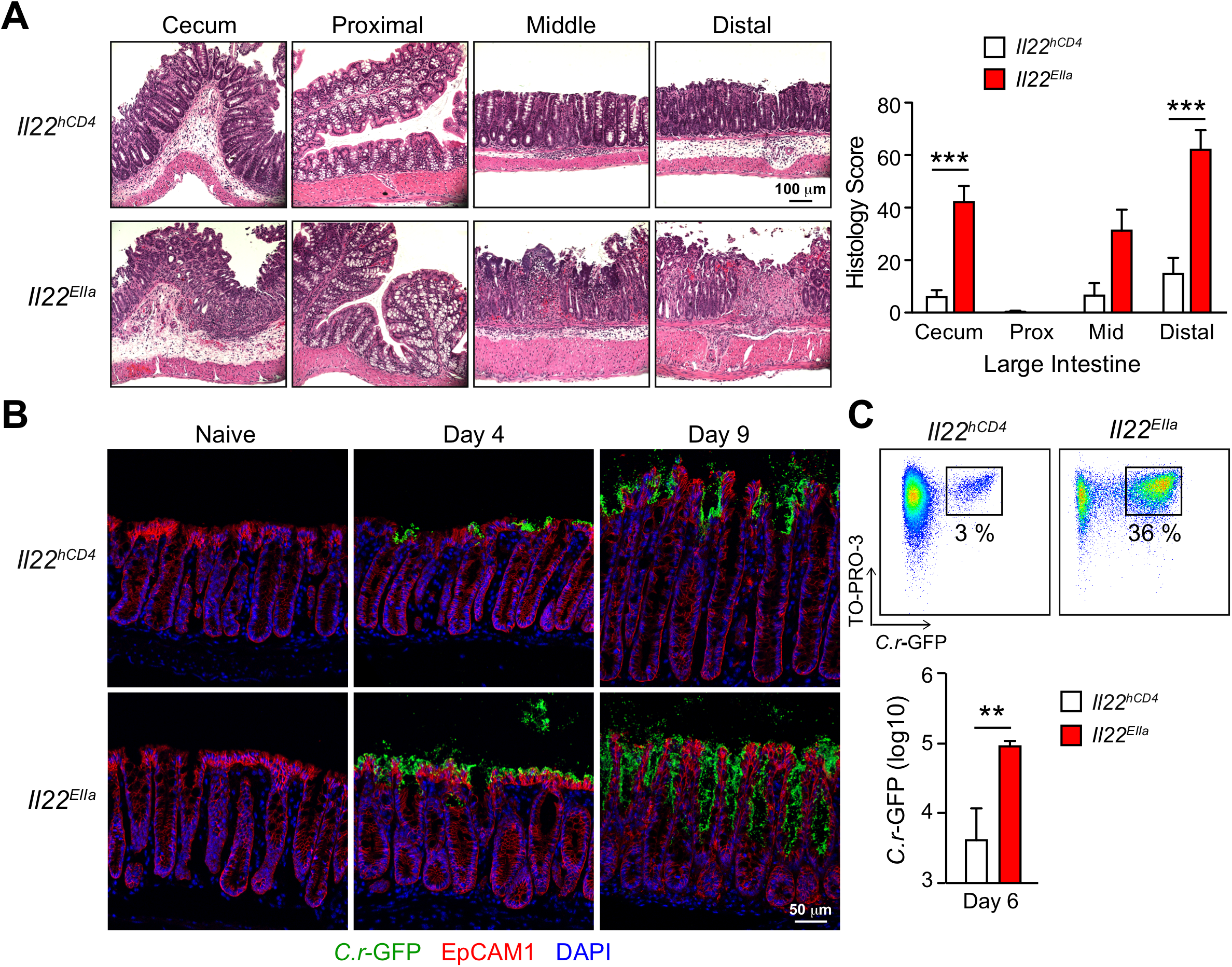
IL-22 protects the colonic crypts from deep bacterial invasion. (A) LI from d8 *C.r*-infected *Il22*^hCD4^ (Cntrl) and *Il22*^EIIa^ (gKO; red) mice was stained with hematoxylin and eosin for blinded histological scoring (Two-way ANOVA, \*\*\**p* <0.001). (B) Colon tissue from naïve and *C.r*-GFP-infected *Il22*^hCD4^ and *Il22*^EIIa^ mice was stained for GFP (green), EpCAM1 (red) and DAPI (blue). (C) *C.r* collected from supernatants of colon IEC preps from d6 *C.r-*GFP-infected Cntrl and gKO mice were stained with TO-PRO-3 and analyzed by flow cytometry in log scale (Mann Whitney, \*\**p* <0.01). 3-5 mice per group, 2 independent experiments.

To define the contributions of IL-22^+^ ILCs and T cells to restraint of the spread of *C.r* across the surface of IECs in *Il22*^EIIa^ gKO mice, parallel studies were performed in *Il22*^Plzf^ and *Il22*^ΔTcell^ cKO mice. Similar to gKO mice and consistent with our bioluminescent studies, *C.r*-infected *Il22*^Plzf^ cKO mice had enhanced bacterial colonization of surface IECs with ∼10-fold higher bacterial burden compared to controls on d4 of infection (**Figures 2E** and **2F** and **Figures 4A** and **4B**). However, our tissue staining and bioluminescent studies show that there were no significant differences in the distribution and bacterial load on d9 of infection, consistent with a dominant influence of IL-22^+^ CD4 T cells by this stage of infection (**Figures 1**, **2E** and **2F,** and **4A** and **4B**). Importantly, and in contrast to *Il22*^EIIa^ gKO mice, there was no significant extension of *C.r* into the crypts of infected *Il22*^Plzf^ cKO mice either prior to death in those mice that were sacrificed due to moribundity or the fraction of mice that survived the innate phase infectious “crisis” (**Figure 4A**). Moreover, histopathologic exam of *Il22*^Plzf^ cKO mice that survived did not show significant differences from controls in colitis scores at the peak of infection (**Figures S2E** and **S3B**). These data reinforced the importance of innate immune cell-derived IL-22 in restraining bacterial proliferation and spread across the superficial epithelium, but suggested that ILC3-derived IL-22 might be inadequate for protection of the crypts.

**Figure 4.**
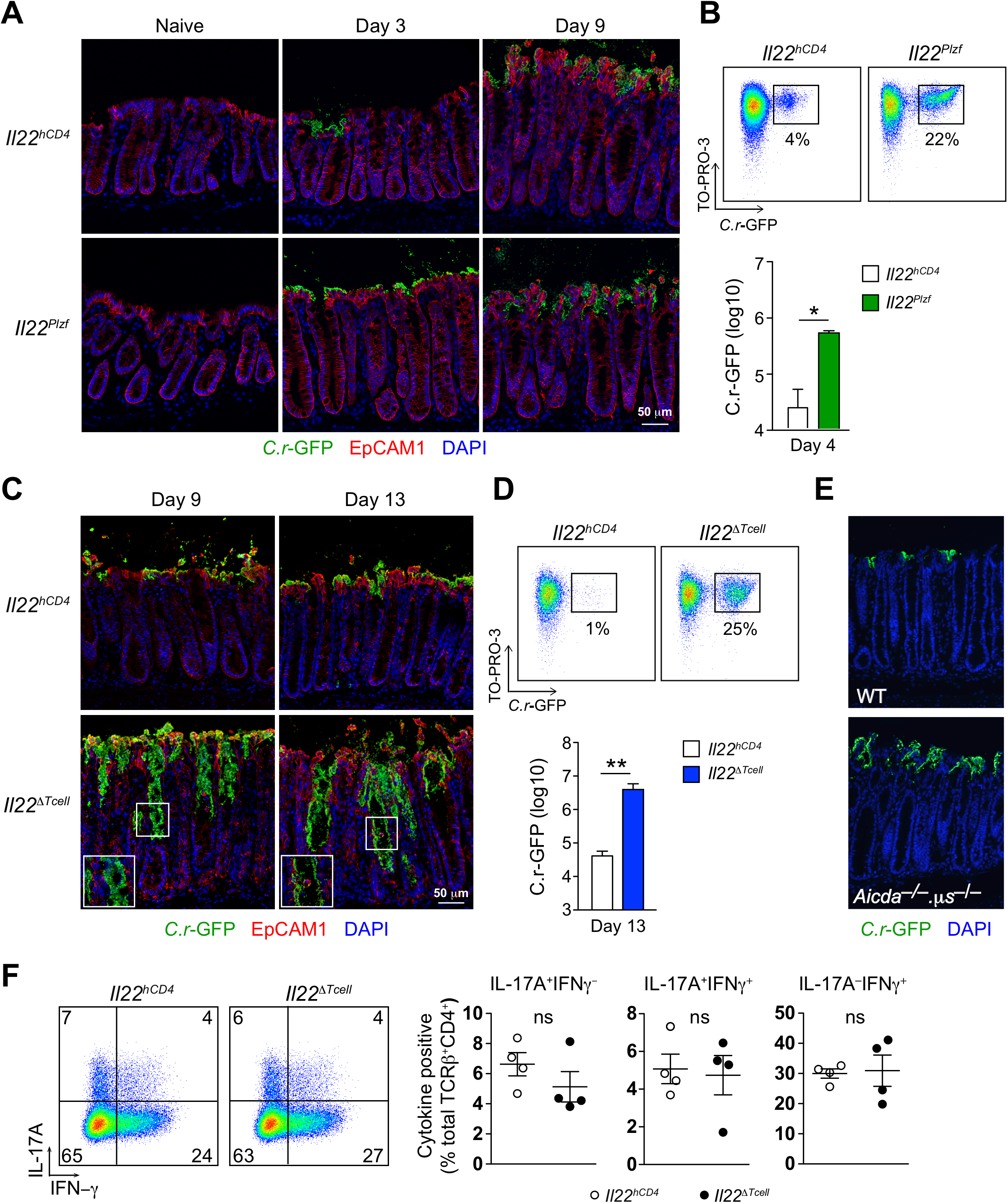
IL-22-expressing innate and adaptive cells protect distinct regions of the colon during *C.r* infection. (A) Colon tissue from naïve and *C.r*-GFP-infected *Il22*^hCD4^ (Cntrl) and *Il22*^Plzf^ (Innate cell cKO) mice was stained for GFP (green), EpCAM1 (red) and DAPI (blue). (B) *C.r* from supernatants of colon IEC preps from d4 *C.r-*GFP-infected *Il22*^hCD4^ and *Il22*^Plzf^ (green) mice was stained with TO-PRO-3 and analyzed by flow cytometry in log scale (Mann Whitney, \**p* <0.05). (C) Colon tissue from d9 and d13 *C.r*-GFP-infected *Il22*^hCD4^ and *Il22*^ΔTcell^ (T cell cKO) mice was stained for GFP (green), EpCAM1 (red) and DAPI (blue). (D) *C.r* from supernatants of colon IEC preps from d13 *C.r-*GFP-infected Cntrl and *Il22*^ΔTcell^ (blue) mice was stained with TO-PRO-3 and analyzed by flow cytometry in log scale (Mann Whitney, \*\**p* <0.01). (E) Colon tissue from d15 *C.r-*GFP-infected WT and *Aicda*^−/−^ □*s*^−/−^ mice was stained for GFP (green) and DAPI (blue). (F) Colon LPLs from *C.r*-infected Cntrl (open) and *Il22*^ΔTcell^ (solid) mice were stimulated with PMA/Ion and Brefeldin A for 4 hrs and then stained with for TCRβ, mCD4, and LIVE/DEAD dye, followed by intracellular staining for IL-17A and IFN□□. ns=not significant, nd=not detected. 3-4 mice per group, 2 independent experiments.

The discrepancy in crypt invasion by *C.r* between *Il22*^EIIa^ gKO mice versus *Il22*^Plzf^ cKO mice suggested that T cell-derived IL-22 might be required for crypt protection. Indeed, this proved to be the case. Strikingly, we found that while during the early course of infection there was no significant difference in bacterial load in control versus *Il22*^ΔTcell^ mice due to an intact ILC response (**Figure 2H**), at later time points, which correlated with the influx of T cells in the colonic LP, *C.r* cells were observed to extend into the colonic crypts (d9; **Figure 4C**), ultimately extending to the bases of crypts by days 12-14 of infection (**Figure 4C**, right panels)—similar to our findings in surviving gKO mice (**Figure 3B,** and **data not shown**). In accord with the widespread extension of *C.r* into crypts, and whole-body imaging data (**Figures 2G** and **2H**), flow cytometric quantitation of *C.r* attachment to IECs was ∼100-fold higher in *Il22*^ΔTcell^ cKO mice compared to controls during this late phase of the response (**Figure 4D**). Furthermore, *C.r*-infected *Il22*^ΔTcell^ cKO mice had significantly worse tissue pathology prior to death (**Figure 2B**), including increased hyperplasia, goblet cell loss, and crypt cell injury in the middle and distal colon compared to *C.r*-infected control mice (**Figures S4A** and **S4B**).

The lack of protection of the crypts in *Il22*^ΔTcell^ cKO mice could not be attributed to altered production of protective antibodies against *C.r*, which are ultimately required for complete clearance of infection (Bry and Brenner, 2004; Maaser et al., 2004; Simmons et al., 2003), as infected mice with normal B cell numbers but deficiency of both Ig class-switching and secreted IgM (*Aicda*^−/−^.*μs*^−/−^) showed protection of crypts that was comparable to control mice (**Figure 4E**). Moreover, total fecal IgG was elevated in *Il22*^ΔTcell^ cKO mice, perhaps reflecting the increased bacterial load (**Figure S4C**). Furthermore, deficient protection of crypts was not due to deficiency of non-IL-22 producing effector CD4 T cells in *Il22*^ΔTcell^ cKO mice, as the number and effector phenotype of these cells isolated from the LP of infected mice did not differ significantly from controls (**Figure 4F**). Thus, IL-22 produced by CD4 T cells are essential for protection of colonic crypts from bacterial invasion; IgG antibodies that are required for the ultimate clearance of *C.r* were not required for this protective function and innate cell-derived IL-22 were unable to protect the crypts. Collectively, these data establish that while ILC3s were sufficient to restrain *C.r* colonization early in infection, only IL-22–producing T cells protected the colonic crypts from bacterial invasion. Thus, IL-22–expressing innate and adaptive immune cells have distinct, specialized roles in the clearance of attaching and effacing enteric pathogens.

### IL-22–producing innate and adaptive immune cells target different subsets of IECs

Because our findings implied a different capacity of IL-22–producing ILC3s and CD4 T cells to activate a protective response in colonic crypts, we reasoned that this might reflect differential IL-22 signaling into IEC subsets. Previous studies have demonstrated that STAT3 signaling is crucial for the protective effects of IL-22 on epithelial cells (Pickert et al., 2009; Sovran et al., 2015; Wittkopf et al., 2015). However, details on which subsets of epithelial cells are activated by IL-22 and from what cellular source are unclear. We therefore surveyed the colonic mucosa of *Il22*^hCD4^ WT and *Il22*^EIIa^ gKO mice for STAT3 activation by immunostaining for phosphorylation of tyrosine residue 705 (pTyr705; pSTAT3) at steady state and over the course of infection. Interestingly, we found that at steady state in naïve control mice, pSTAT3 was undetectable both in colonic IECs or cells in the LP (**Figures 5A** and **5C**); i.e., no baseline activation of STAT3 was evident. During the innate phase of *C.r* infection (d4), low-intermediate intensity pSTAT3 activation (pSTAT3^dim^) was detected in a distribution limited to the nuclei of surface IECs (i.e., those facing the intestinal lumen or lining the mouth of crypts) and in many immune cells in the LP of control mice (**Figures 5A** and **5B**, see insets). Global deficiency of IL-22 completely eliminated detectable pSTAT3 in IECs, but was preserved in immune cells populating the lamina propria, indicating that IL-22 is non-redundant in its activation of IECs during the innate phase of *C.r* infection, whereas other STAT3-activating cytokines signal into LP-resident immune cells. It is notable that the vast majority of superficial IECs were pSTAT3-positive, despite the very focal attachment of small colonies of *C.r* at this time point (**Figure 3B**). This suggests that direct effects of *C.r* due to bacterial attachment and injection of bacterial proteins into IECs neither contribute to, nor interfere with, IEC STAT3 activation driven by IL-22.

**Figure 5.**
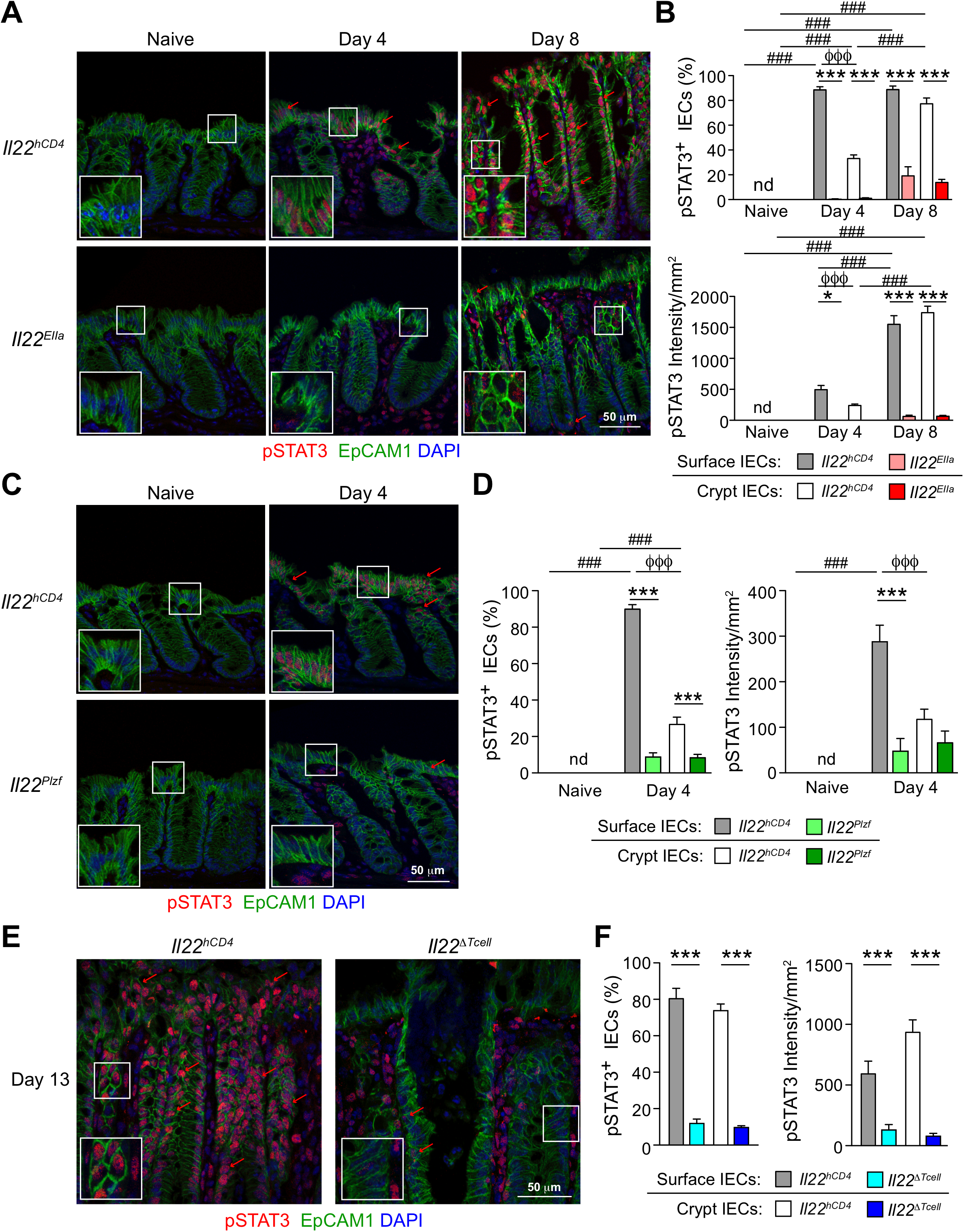
IL-22^+^ T cells induce robust and prolonged STAT3 activation. (A, C, E) Colons from (A) naïve, d4 and d8 *C.r*-infected *Il22*^hCD4^ (Cntrl) and *Il22*^EIIa^ (gKO) mice, and (C) naïve and d4 Cntrl and *Il22*^Plzf^ (Innate cell cKO) mice, and (E) d13 Cntrl and *Il22*^ΔTcell^ (T cell cKO) mice were stained for EpCAM1 (green), pSTAT3 (red) and DAPI (blue). Red arrows depict pSTAT3^+^ IECs. (B, D, F) Quantitation of percent pSTAT3^+^ cells and intensity of pSTAT3 staining in surface and crypt IECs from (B) naïve, d4 and d8 Cntrl and gKO mice, and (D) naïve and d4 *C.r*-infected Cntrl and *Il22*^Plzf^ mice and (F) d13 Cntrl and *Il22*^ΔTcell^ mice. (B, D) Two-way ANOVA with Bonferroni posttests; ^###^*p* <0.001 comparing different time points. \**p* <0.05 and \*\*\**p* <0.001 comparing WT and cKO mice and ^ϕϕϕ^*p* <0.001 comparing surface and crypt IECs. (F) One-way ANOVA with post-hoc Tukey tests; \*\**p* <0.01 and \*\*\**p* <0.001 comparing WT and cKO mice. nd=not detected. 4-5 mice per group, 2 independent experiments.

In contrast to superficial IECs, most IECs lining the crypts were pSTAT3-negative or showed minimal pSTAT3 during the innate phase of infection (d4). This changed dramatically with the influx of IL-22^+^ CD4 T cells (d8) (**Figures 5A** and **5B**). While there was no significant difference in the number of pSTAT3-positive surface IECs, the average intensity of staining increased ∼3-fold, reflecting higher amplitude pSTAT3 signaling (pSTAT3^bright^). Additionally, IECs now became pSTAT3-bright at all levels of the crypts, with comparable frequencies and staining intensities to those of surface IECs. Strikingly, STAT3 activation was virtually ablated in surface and crypt IECs in *Il22*^EIIa^ gKO mice compared to controls, indicating that IL-22 is also indispensable for STAT3 activation of crypt cells during *C.r* infection. Because the crypts of WT mice are devoid of invading bacteria, this further reinforces the notion that direct attachment of *C.r* is not required for local STAT activation in IECs. Also, as noted above, no discernable decrement in the frequency or intensity of pSTAT3 was evident in immune cells within the LP consequent to loss of IL-22 during the late, adaptive phase of infection. Thus, whereas IL-22 is indispensable for activation of IECs, other STAT3-activating cytokines act on immune cells in the involved mucosa, consistent with their expression of other STAT3-activating cytokine receptors (e.g., IL-6R or IL-23), but not IL-22R.

To extend these findings, we examined IL-22–dependent activation of STAT3 in IECs contingent on the source of IL-22, whether from ILC3s or T cells. During the early stages of infection when ILC3-derived IL-22 is dominant in limiting *C.r* colonization of the luminal surface, STAT3 activation was significantly diminished in surface IECs of *C.r*-infected *Il22*^Plzf^ cKO mice compared to *C.r*-infected control mice (**Figures 5C** and **5D**, see insets). Importantly, during the late phase of *C.r* infection when T cell-derived IL-22 is required for crypt protection, both the frequency and intensity of pSTAT3 positivity was markedly reduced in IECs of *C.r*-infected *Il22*^ΔTcell^ cKO mice compared to controls (**Figures 5E** and **5F**, see insets). Remarkably, deficiency of T cell-derived IL-22 also ablated STAT3 activation in surface IECs, despite the persistence and sustained competency of ILC3s to produce IL-22 during this period (**Figure 1**). Thus, ILC3-derived IL-22 was no longer sufficient to induce STAT3 activation in surface IECs. Instead, and akin to crypt IECS, late in infection the surface IECs were strictly dependent on T cell-derived IL-22 for STAT3 activation and thus protective responses.

Taken together with our previous findings, these data establish that IL-22 produced by ILCs is the principal cytokine driving IEC STAT3 activation that is necessary for restraint of *C.r* colonization and host survival during the early course of infection, but its effectiveness is limited to surface IECs and is extinguished as the infection progresses. As the infection progresses, T cell-derived IL-22 is indispensable for driving STAT3 activation that underpins resistance of crypt IECs to *C.r* invasion and for sustaining and amplifying STAT3 activation of surface IECs when ILC3s no longer support this function. Thus, the non-redundant function of IL-22 in host protection against attaching/effacing bacteria reflects the unique ability of this cytokine to activate STAT3 in IECs, and CD4 T cells are indispensable for protection of both the colonic crypts and the colonic surface barrier as *C.r* infection progresses.

### T cell-derived IL-22 promotes a shift in IEC functional programming to protect intestinal crypts

All subsets of IECs arise from intestinal stem cells (ISCs) that are sequestered from the intestinal lumen—and potential pathogens—in the base of intestinal crypts (Barker et al., 2007; Chang and Leblond, 1971; Hua et al., 2012; van der Flier and Clevers, 2009). The differentiation and specialization of IECs in the colon occur as progeny of ISCs divide and transit along the crypt-surface axis, giving rise to absorptive enterocytes (ECs), which are the major surface IEC, and secretory IECs, including goblet cells, tuft cells, enteroendocrine cells (EECs), and, in the colon, deep secretory cells (DSCs, or Paneth-like cells) that appear to share ISC-supportive functions similar to that of Paneth cells in the small intestine (Rothenberg et al., 2012; Sasaki et al., 2016). Because our findings identified a unique role for IL-22-producing T cells in activating STAT3 signaling in crypt IECs—including those residing in the IEC “incubator” at the base of crypts—during *C.r* infection, we sought to understand how T cell-derived IL-22 might reprogram developing IECs to protect the crypts.

Genes that were differentially expressed contingent on T cell-derived IL-22 were identified by RNA-seq analysis performed on three subpopulations of FACS-sorted IECs from the mid-distal colons of naïve (d0) and d9 *C.r*-infected control (*Il22*^hCD4^; Cntrl) and T cell cKO (*Il22*^ΔTcell^) mice. Subpopulations were defined on the basis of differential cell size/complexity and expression of EpCAM1 and CEACAM1: Small crypt (SC) cells (EpCAM1^+^ CEACAM1^lo^ FSC^lo^ SSC^lo^); large crypt (LC) cells (EpCAM1^+^ CEACAM1^int^ FSC^hi^ SSC^int^); and superficial, or surface, cells (Srf) (EpCAM1^+^ CEACAM1^hi^ FSC^hi^ SSC^hi^) (**Figures 6A-6D**); which correlated with lower crypt cells, upper crypt cells, and surface cells, respectively, based on correlative gene expression data from FACS-sort/RNA-seq and laser capture microdissection (LCM)/RT-PCR analyses (**Figures S5A-S5D).** At the peak of *C.r* infection (d9), 739 differentially expressed genes (DEGs) were identified in colonic IECs from control versus *Il22*^ΔTcell^ cKO mice (**Figures 6D-6F** and **Figures S5E-S5G**). Although the diversity of genes induced by T cell-derived IL-22 was considerable, several findings were notable.

**Figure 6.**
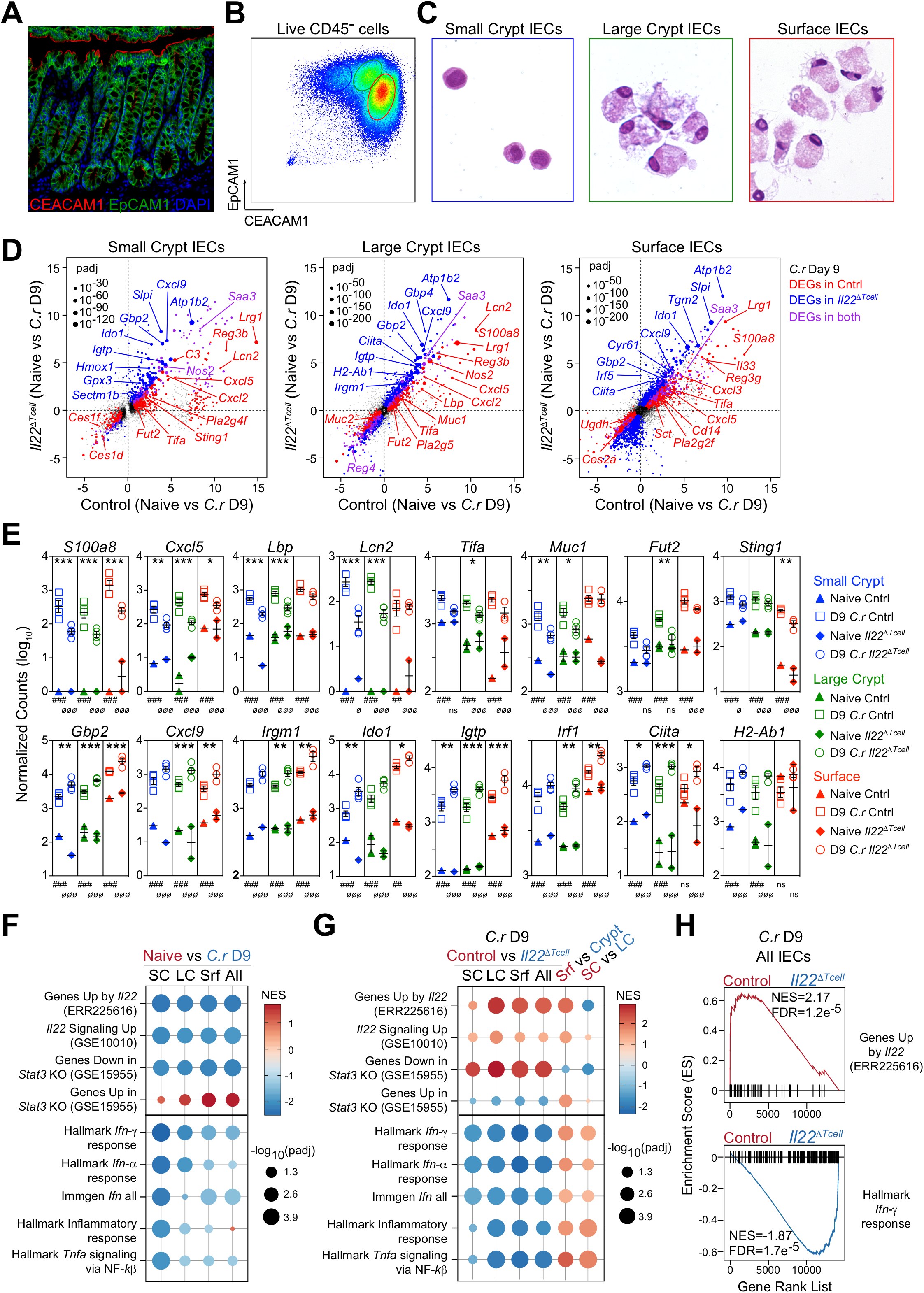
IL-22^+^ T cells upregulate host defense genes and repress IFNDγ-induced genes. (A) Colon tissue from naïve Bl/6 mice was stained for EpCAM1 (green), CEACAM1 (red) and DAPI (blue). (B) IECs from naïve BL/6 mice were stained for EpCAM1, CEACAM1, Live/Dead (L/D) dye and CD45 and analyzed by flow cytometry or (C) sorted into small crypt (SC; EpCAM1^+^ CEACAM1^lo^ CD45^−^ L/D dye^−^ FSC^lo^ SSC^lo^; blue), large crypt (LC; EpCAM1^+^ CEACAM1^int^ CD45^−^ L/D dye^−^ FSC^hi^ SSC^int^; green) and surface IECs (Srf; EpCAM1^+^ CEACAM1^hi^ CD45^−^ L/D dye^−^ FSC^hi^ SSC^hi^; red) and then cytospun and stained with hematoxylin and eosin. (D-H) RNA-seq was performed on sorted SC, LC and Srf IECs from mid/distal colons of naïve and *C.r* D9 *Il22*^hCD4^ (Cntrl) and *Il22*^ΔTcell^ (T cell cKO) mice. (D) Two-way scatter plots comparing DEGs in SC, LC and SF IECs from naïve vs *C.r* D9 Cntrl mice (red) and naïve vs *C.r* D9 *Il22*^ΔTcell^ mice (blue). Purple indicates DEGs in both (*p*adj <0.05; colored dots). (E) Count plots of DE host defense genes in SC (blue), LC (green) and Srf (red) IECs from D9 *C.r* Cntrl (solid) and D9 *C.r Il22*^ΔTcell^ (open) mice. Counts were normalized by library size. \**p*adj <0.1, \*\**p*adj <0.01, \*\*\**p*adj <0.001. (F) Dot plot of gene set enrichment analysis (GSEA) of IL-22, IFN*α*□ IFN*γ*, TNF and Inflammatory pathways in SC, LC, Srf and pooled IECs (All) from Naïve (brick) vs D9 *C.r*-infected Cntrl (azure) mice. (G) Dot plot of GSEA of IL-22, IFN*α* IFN*γ*, TNF and Inflammatory pathways in SC, LC, Srf and pooled IECs (All) from D9 *C.r*-infected Cntrl (brick) vs D9 *C.r*-infected *Il22*^ΔTcell^ (azure) mice, Srf (brick) vs pooled Crypt (azure), and SC (brick) vs LC (azure). (H) Bar code plots of GSEA of IL-22 and IFN*γ* pathways in pooled IECs (All) from *C.r* D9 Cntrl (brick) versus *C.r* D9 *Il22*^ΔTcell^ (azure) mice. NES, normal enrichment score; FDR, false discovery rate. 2-3 mice per sample, 1-2 independent experiments per naïve group and 3-4 independent experiments per infected group.

Transcripts of multiple genes involved in host defense were significantly up-regulated by T cell-derived IL-22, whether predominantly in surface IECs (e.g., *Sting1*), crypt IECs (e.g., *Lcn2*, *Lbp, Muc1*) or all IEC subsets (e.g., *S100a8*, *Lrg1, Tac1*) (**Figures 6D** and **6E** and **Figures S5E-S5G**). The striking induction in crypt IECs of transcripts that encode lipocalin 2, a principal sequestrator of iron-binding siderophores expressed by pathogenic *E. coli* and *C.r* (T. Berger et al., 2006; Goetz et al., 2002), and S100a8, a component of the metal-chelator calprotectin (Brandtzaeg et al., 1995; Clohessy and Golden, 1995), is consistent with an important role of these antimicrobial peptides (AMPs) in defense of the colonic crypts. Moreover, IL-22–producing T cells up-regulated several phospholipase A2 (PLA2) genes that encode for phospholipid-hydrolyzing enzymes that possess bactericidal activity and contribute to IL-22/STAT3-dependent host defenses (Harwig et al., 1995; Okita et al., 2016; Wittkopf et al., 2015; Yamamoto et al., 2015) (**Figure 6D** and **Figures S5E-S5G**, **Figures 7B** and **7C** and **Figure S6**). Notably, although transcripts encoding the Reg family antimicrobial peptides, Reg3*β* and Reg3*γ*—previously implicated as important IL-22-dependent AMPs in *C.r* infection (Zheng et al., 2008)—were induced by T cell-derived IL-22 at sites of *C.r* colonization in the mid/distal colon, their expression was found to be significantly higher in the proximal colon, which is not colonized by *C.r* during infection (Basu et al., 2012; Wiles et al., 2004) (**Figure 6D** and **Figures S5E-S5G**, and data not shown). Up-regulation in crypt IECs of transcripts encoding the LPS-binding protein (*Lbp*), a key factor in enhanced recognition of the Gram-negative bacterial cell wall component by TLR2 and TLR4 (Medzhitov et al., 1997; Poltorak et al., 1998; Pugin et al., 1993; Schletter et al., 1995), suggests that potentiation of recognition of this pathogen-associated molecular pattern (PAMP) by crypt IECs may contribute to crypt defense in *C.r* infection.

**Figure 7.**
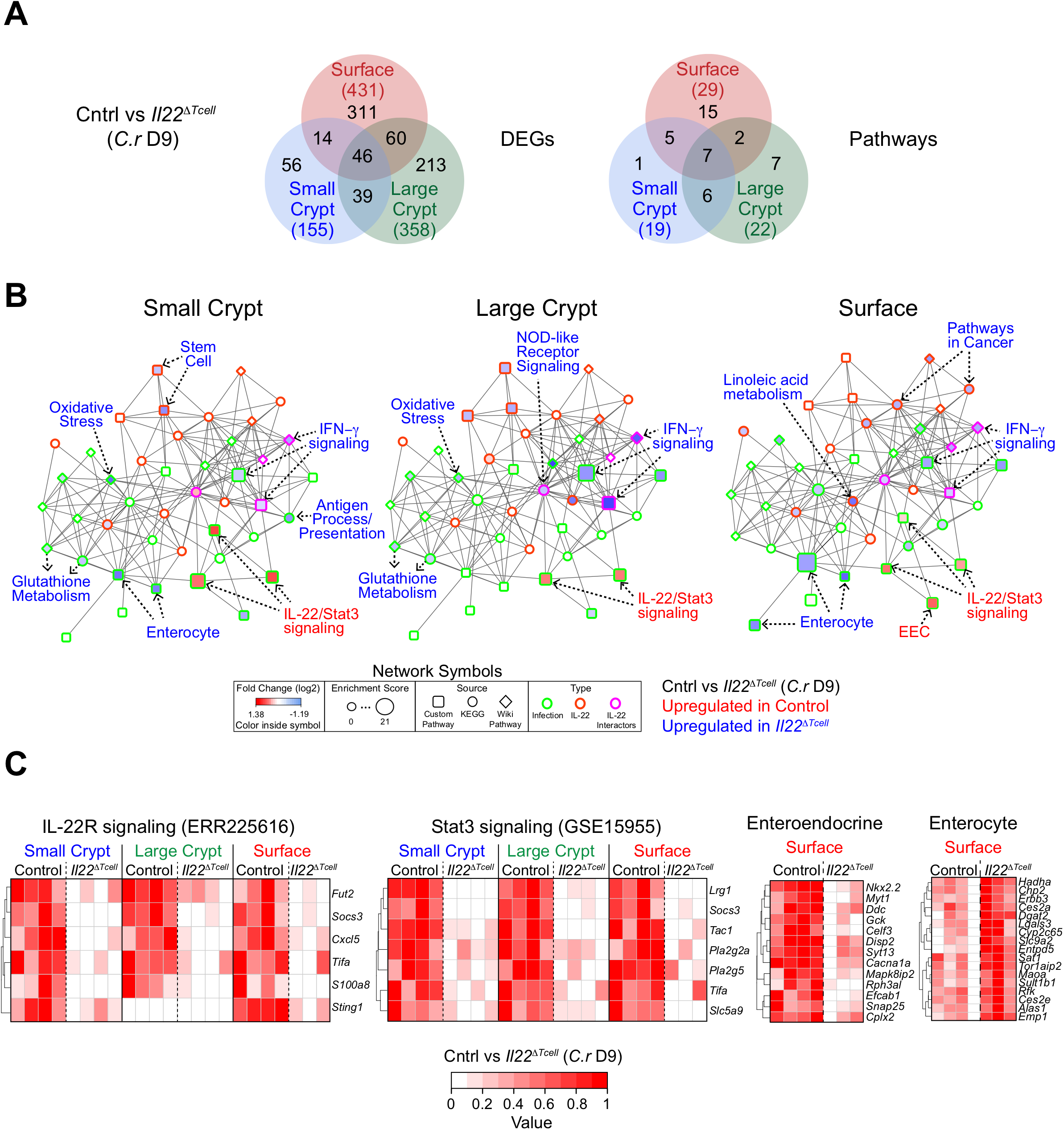
T cell-derived IL-22 promotes a shift in IEC functional programming to protect intestinal crypts. (A-C) RNA-seq was performed on sorted SC, LC and Srf IECs from mid/distal colons of *C.r* D9 *Il22*^hCD4^ (Cntrl) and *Il22*^ΔTcell^ (T cell cKO) mice. (A) Venn diagram of DEGs and Pathways in SC (red), LC (green) and Srf (red) IECs from *C.r* D9 Cntrl and *Il22*^ΔTcell^ mice. Numbers in parenthesis depict total number of DEGs or Pathways per IEC subset. (B) Pathway analysis in SC, LC and Srf IECs from *C.r* D9 Cntrl and *Il22*^ΔTcell^ mice. Color inside the symbol depicts fold change (log_2_, pathway up in Cntrl (red), pathway down in Cntrl (blue)), size of the symbol depicts enrichment score, shape of the symbol depicts source (Custom Pathway (square), KEGG (circle), Wiki Pathway (diamond)), and border color depicts type of interaction (Infection (green), IL-22 (orange), IL-22 interactors (pink)). (C) Heatmaps of the top DEGs in IL-22R, STAT3 and custom IEC pathways. 2-3 mice per sample and 3-4 independent experiments per group.

T cell-derived IL-22 also played a significant role in the induction of neutrophil-attractant chemokine expression and a shift in mucin production and modification by IECs that contribute to host defense during *C.r* infection (Aujla et al., 2008; Bergstrom et al., 2008; Hopkins et al., 2019; S. C. Liang et al., 2010; Lindén et al., 2008) (**Figures 6D** and **6E** and **Figures S5E-S5G**). Transcripts for *Cxcl1*, *Cxcl2* and *Cxcl5*, which are recognized by neutrophils via CXCR2, were induced in both surface and crypt IECs, consistent with a central role for IL-22 signaling into IECs to initiate the recruitment of neutrophils into the infected mucosa and infected crypts, where they contribute to clearance of *C.r* (Kamada et al., 2015). Previous studies have identified an IL-22–induced shift in mucin production and its altered mucin fucosylation by IECs (Pham et al., 2014; Sugimoto et al., 2008; Turner et al., 2013), which we find are dependent on IL-22-expressing T cells. Thus, in addition to its role in amplifying antimicrobial recognition and production of AMPs, T cell-derived IL-22 may coordinate the recruitment of neutrophils to the infected colonic lamina propria and lumen, and a shift in mucin production from Muc2 to Muc1 (**Figure 6D** and **6E**, **Figures S5E-S5G** and **Figure S4D**), as well as altered mucin fucosylation, which may deprive *C.r* of an important energy source (Pham et al., 2014).

In addition to genes that were induced, genes normally repressed by T cell-derived IL-22 were a major component of the DEG profile in each IEC subset (**Figures 6D-6H** and **Figures S5E-S5G**). Notable from a combined gene set, network and pathway analysis (GNPA) was prominence of genes induced by TNF*α*and IFN*γ* that were significantly increased in absence of T cell-derived IL-22 (**Figures 7A** and **7B** and **Figure S6**). This included genes of the antigen processing and presentation pathway, particularly MHC class I and II genes, which reflected up-regulation of the gene encoding the central transactivator of this pathway, *Ciita* (Martin et al., 1997; Steimle et al., 1993). Proinflammatory genes activated by interferon signaling were also repressed by IL-22 signaling (**Figures 6D-6H, Figures 7A** and **7B**, **Figures S5E-S5G** and **Figures S6A** and **S6B**), including the IFN*γ*-dependent chemokine transcripts *Cxcl9* and *Cxcl10*, which act as chemoattractants for immune cells recruited in type I responses, including Th1 cells (Loetscher et al., 1996; Luster and Ravetch, 1987). Because IFN*γ* is required for goblet cell loss and IEC proliferation during *C.r* infection (Chan et al., 2013), heightened IFN*γ* responses due to the deficit of IL-22 signaling may have contributed to the enhanced goblet cell hypoplasia and IEC damage, as well as a trend towards increased crypt hyperplasia observed in *Il22*^ΔTcell^ cKO mice (**Figures S4A** and **S4B**). Moreover, DEGs repressed by IL-22 were characteristic of absorptive enterocytes (ECs; e.g., *Ces2c*, *Cyp3a13*, *Ubd*, *Ugdh*, *Noct*) (**Figure 6D**, **7B** and **7C** and **Figures S5E-S5G** and **S6**), as reflected in the enhanced enterocyte signature in the GNPA analysis (**Figure 7B** and **7C** and **Figure S6**). Strikingly, many genes characteristic of mature ECs were enriched in Srf IECs from *C.r*-infected *Il22*^ΔTcell^ cKO mice compared to infected controls, suggesting that T cell-derived IL-22 normally acts to repress maturation of ECs driven by IFN*γ*-driven hyperproliferation during infection, perhaps as a measure to deprive *C.r* of its cellular host for attachment and colonization. This was contrasted by enhancement of the enteroendocrine cell gene signature in Srf IECs (e.g., *Tac1*, *Adgrl1*, *Celf3*, *Myt1*, *Sct*) (**Figure 6D**, **7B** and **7C** and **Figures S5E-S5G** and **S6**), implicating an important regulatory role for T cell-derived IL-22 in programming EEC differentiation and/or function to alter the local expression of EEC-derived hormones. Collectively, these findings indicate that, in addition to shifting the type of mucus produced by IECs and enhancing the expression of a select set of AMPs and chemokines, IL-22 signaling provided by T cells plays an important role in modulating the development of IECs that may restrain bacterial invasion of the crypts—whether by promoting STAT3 activation to induce gene expression or repress aspects of interferon and TNF signaling.

## Discussion

In this study we define a non-redundant role for IL-22-producing T cells in antibacterial defense of colonic crypts. Our findings address a central, unresolved issue regarding the coordination of innate and adaptive immunity concerning the specialization of ILCs and CD4 T cells. Since the discovery of ILC subsets and the emergent appreciation of their functional parallels and redundancy with T-cell subsets (Bando and Colonna, 2016; Huntington et al., 2016; Spits et al., 2013), it has been unclear what functions are unique to each immune cell population. Here we find that, despite their critical role in restraining bacterial colonization over the early course of enteropathogenic bacterial infection, ILC3s—and other IL-22-producing innate immune cells—induce relatively weak STAT3 signaling that is limited to surface IECs; they prove unable to activate IL-22-dependent STAT3 signaling in IECs of the colonic crypts. Rather, it is T cells that are uniquely charged with delivery of IL-22 to crypt IECs and to surface IECs as infection progresses, inducing more robust, sustained STAT3 signaling in both IEC populations that is required to activate programs essential for host defense against bacterial invasion.

Although the mechanisms by which IL-22–producing T cells achieve heightened activation of IECs are not yet fully defined, it would appear that a major, if not sole, contributor is the geography of immune-cell positioning relative to the intestinal epithelium. Whereas we find that most ILC3s are sequestered in mucosal lymphoid tissues and fail to increase their local numbers throughout infection (Ahlfors et al., 2014; Gasteiger et al., 2015), CD4 T cells generated in response to infection become the major population of IL-22-positive cells in the inflamed mucosa and are positioned immediately adjacent to the epithelium they are charged with protecting, particularly the crypts. Here they are uniquely able to activate IL-22/STAT3 signaling into crypt IECs, and also become the sole source for sustained activation of surface IECs as the quality of these cells is altered by a shift in IEC maturation during infection. It will be important to determine whether this is due to production of increased local concentrations of IL-22, directed delivery of IL-22 to IECs in the context of MHC class II-mediated non-classical antigen presentation, delivery of co-signals that amplify IL-22-mediated STAT3 activation, or a combination of these. Irrespective of mechanism, T cells would appear to deliver IL-22 to IECs on-site, whereas ILCs must deliver IL-22 long-range.

The host-protective effects consequent to actions of T cell-derived IL-22 on IECs are diverse and non-redundant. Consistent with a previous study showing that STAT3-activating IL-22 and not IL-6 participates in IEC activation during DSS-induced colitis (Pickert et al., 2009), we find that IL-22 induces detectable STAT3 activation during each phase of *C.r* infection, reflective of this cytokine’s critical role in antibacterial host defense (Basu et al., 2012; Sonnenberg et al., 2011b; Zheng et al., 2008). Major actions of T cell-derived IL-22 evident from our RNA-seq analyses include the induction of antimicrobial peptide (AMPs) and neutrophil-recruiting chemokines (Aujla et al., 2008; Boniface et al., 2005; S. C. Liang et al., 2010; 2006; Okita et al., 2016; Wittkopf et al., 2015; Wolk et al., 2006; Yamamoto et al., 2015; Zheng et al., 2008), alteration of mucin production and fucosylation (Pham et al., 2014; Sugimoto et al., 2008), and enhancement of other defenses that restrain bacterial growth (e.g., induction of *Sting1* and *Lbp*) (Aden et al., 2018; Wolk et al., 2006; 2007). Based on analysis of whole colon tissue from global IL-22 deficient mice, it has been proposed that host protection mediated by IL-22 is due to upregulation of the Reg3 family of antimicrobial peptides (AMPs) (Zheng et al., 2008), particularly Reg3D, which, unlike Reg3D (Cash et al., 2006; Pham et al., 2014), has anti-microbial actions against Gram-negative bacteria (Miki et al., 2012; Stelter et al., 2011). Although our gene expression studies are consistent with an important role for T cell-derived IL-22 in the induction of AMPs, we find that, while transcripts encoding *Reg3b* and *Reg3g* are up-regulated in areas of *C.r* colonization—the mid- and distal colon—their expression is far greater in the proximal colon, which is not colonized during *C.r* infection (Basu et al., 2012; Wiles et al., 2004). This is in contrast to the upregulation of the *S100a* family of AMPs and *Lcn2*, which occurs in the areas colonized by *C.r*. It is unclear if this reflects a particularly potent effect of IL-22–induced Reg3D in colonization resistance or rather a limited role for this AMP in protection against *C.r*, as has been shown for RegDD (Pham et al., 2014). It will be important to determine which, if any, of the AMPs induced by T cell-derived IL-22 may be important for defense of the crypts.

In a previous study, invasion of colonic crypts by *C.r* was observed in *Cxcr2*-deficient mice, which have a profound defect in recruitment of neutrophils to the infected colon (Spehlmann et al., 2009). Our finding that T cell-derived IL-22 upregulates in IECs transcripts for several CXCL chemokines that are ligands of CXCR2 (e.g., *Cxcl1*, *Cxcl2*, and *Cxcl5*) indicates a mechanism by which neutrophil recruitment during *C.r* infection may be initiated, and, in view of the important role for neutrophils in eradicating luminal bacteria (Kamada et al., 2015), indicates that this pathway controls a central mechanism for the antibacterial defense of crypts. Because neutrophils are themselves an important source of CXCL2 (Li et al., 2016), the finding of IL-22-dependent induction of CXCR2-binding chemokines suggests a feed-forward mechanism whereby initial recruitment of neutrophils to the infected mucosa is initiated by T cell-derived IL-22 activation of IECs that may be amplified by incoming neutrophils. Interestingly, in preliminary studies we find that the expression of *Cxcl5* is limited to IECs (Cai et al., unpublished observation), raising the possibility that this neutrophil chemoattractant may be important in directing infiltrating neutrophils to sites of bacterial attachment on IECs. This will require further study. In any case, our findings identify a novel link between T cell-derived IL-22 and neutrophil recruitment that may aid in limiting *C.r* invasion of crypts. Interestingly, however, despite the important role for *C.r*-specific IgG responses in the ultimate clearance of infection (Bry and Brenner, 2004; Maaser et al., 2004)—thought to be due in part to antibody-dependent opsonization of the bacterium to enhance neutrophil-mediated phagocytosis (Kamada et al., 2015)—it was surprising that no major role for antibody-dependent protection of the crypts was found, despite its essential role in pathogen clearance. Thus, the actions of neutrophils in defense of colonic crypts appear to be adequate without requirement for IgG-mediated bacterial opsonization, whether via phagocytosis-dependent or - independent mechanisms, or both.

Another major action of T cell-derived IL-22 was its tempering of pro-inflammatory and epithelial differentiation effects on IECs exerted by TNF and interferon signaling. The actions of IFN*γ* signaling on IECs, in particular, have been shown to result in the acceleration of IEC proliferation that characterizes crypt hyperplasia and goblet cell loss that are thought to protect the crypts from bacterial incursion (Chan et al., 2013). In essence, the hyperproliferation of IECs driven by IFN*γ*may be viewed as a mechanism to further distance ISCs in the crypt bases from invading pathogens to protect them from both physical and metabolic insults (Kaiko et al., 2016; Y. Liang et al., 2017; Matsuki et al., 2013; Okada et al., 2013). However, we find that, in the absence of T-cell production of IL-22, the crypts are not protected despite unopposed actions of IFN signaling that result in increased goblet cell loss and crypt hyperplasia. Thus, the alterations in IEC developmental programming induced by IFN*γ* signaling are inadequate without coordinate actions of IL-22 delivered by T cells; IFN*γ* and IL-22 must cooperate in defense of the crypts as deficiency of either leads to bacterial invasion.

Our results raise the intriguing possibility that, in addition to its other actions on IECs, IFND signaling is required for induction of antigen processing and presentation by IECs as a means to recruit more potent, protective IL-22 signaling from Th17 and Th22 cells. Although deficiency of IL-22 resulted in enhanced IEC expression of *Ciita* and thus major components of the antigen processing and presentation pathway, MHC class II expression was nevertheless robust in infected controls (**Figure S4E**), implying a role for IECs as non-conventional antigen-presenting cells (APCs) to directly recruit T cell responses. It was recently reported that antigen presentation by Lgr5^+^ ISCs in the small intestine elicits IL-10 from Foxp3^+^ Treg cells that sustains homeostatic ISC self-renewal (Biton et al., 2018). It was further suggested that antigen presentation by Lgr5^+^ ISCs to effector T cells responding to intestinal infections may recruit T cell cytokines that shift ISC programming from homeostatic self-renewal to host-defensive IEC differentiation. Consistent with this hypothesis—and extending it—we find that T cell-derived IL-22, also a member of the IL-10 cytokine family, drives strong STAT3 activation in *all* IECs, not just ISCs, thereby restraining IFN*γ*-induced IEC differentiation while promoting antimicrobial defense. Notably, we detected no STAT3 activation in colonic IECs at steady state, including ISCs, suggesting that, in contrast to the small intestine, neither IL-10 nor IL-22 has homeostatic actions on IECs in the colon. In any case, in its non-redundant role to defend colonic crypts from bacterial invasion, T cell-derived IL-22, like its family member IL-10, would appear to play an essential role in the STAT3-dependent maintenance of intestinal stem cells as a means to insure restitution of the epithelial barrier and preservation of mucosal integrity. Going forward, it will be important to determine whether recognition of antigen presented on IECs underlies the basis for the unique ability of IL-22-producing T cells, but not innate immune cells, to activate crypt IECs for antimicrobial defense.

## Lead Contact and Materials Availability

Further information and requests for resources and reagents should be directed to and will be fulfilled by the Lead Contact, Casey T. Weaver. The mouse lines obtained from other laboratories are described below and may require a Material Transfer Agreement (MTA) with the providing scientists. *Il22^hCD4^* mice generated in this study are available from our laboratory, also with an MTA.

## Experimental Model and Subject Details

### Mice

*Il22*^hCD4.fl^ reporter/cKO mice were generated by targeting an IRES and truncated hCD4 gene into the fifth exon (between the stop codon and 3’ untranslated region), and loxP sites that flanked the entire *Il22* gene (see Method Details below). B6 Albino (CRL 022) and CD-1 (CRL 493) used for generating chimeric mice were purchased from Charles River Laboratories (CRL). C57BL/6 (WT; JAX 000664), *Actb-Flpe* (JAX 003800), *EIIa*-cre (JAX 003724), *Plzf*-cre (JAX 024529) and *mCd4-*cre (JAX 022071) were purchased from Jackson Laboratory (JAX). *Rorc*^EGFP^ mice were kindly provided by Dr. Gerard Eberl (Lochner et al., 2008) (Instit Pasteur, France). *Aicda*^−/−^.*Ds*^−/−^ mice were kindly provided by Dr. Frances E. Lund (UAB). In most experiments, littermates were used as controls and experimental animals were co-caged in groups of 2-7 mice. Both sexes were used per experimental group whenever possible. All mouse strains were bred and maintained at UAB in accordance with IACUC guidelines.

### Citrobacter rodentium Infections

*Citrobacter rodentium (C.r)* strain, DBS100 (ATCC 51459) was used for all survival and kinetics experiments. For whole-body imaging experiments, the bioluminescent *C.r* strain ICC180 (derived from DBS100) was used (Wiles et al., 2006) (generously provided by Drs. Gad Frankel and Siouxsie Wiles, Imperial College London). Animals were imaged for bioluminesence using an IVIS-100 Imaging System (Xenogen). For flow cytometry analysis and to track *C.r in situ*, a strain of *C.r* expressing GFP (derived from DBS100) was used (Bergstrom et al., 2010) (kindly provided by Dr. Bruce A. Vallance). A fresh, single colony was grown in 5 ml LB overnight at 37**°**C with agitation for 12-14 hrs. Next day, 1 ml of overnight culture was added to 250 ml LB, incubated at 37**°**C with agitation for 5-6 hrs and then stopped when OD600 reached 1.0. Bacteria was pelleted at 4**°**C, 3000 rpm for 15 minutes and then resuspended in 5 ml sterile 1x PBS. Mice were inoculated with 1–2×10^9^ cfu in a total volume of 100 µl of PBS via gastric gavage.

## Method Details

### Generation of *Il22*^hCD4^ reporter/conditional knockout mice

The BAC clone RP24-227B3, which contains the *Il22* gene and *Iltifb* (*Il22* pseudogene) was purchased from Children’s Hospital Oakland Research Institute (CHORI). Briefly, a targeting cassette containing an EMCV IRES, neomycin^R^ cassette, truncated hCD4 gene (pMACS 4-IRES.II; Miltenyi Biotec) and 3’ loxP site flanked with arms of homology to exon 5 of the *Il22* gene was recombineered into RP24-227B3 (see Figure S1A). First, 5’ arm of homology to exon 5 of the *Il22* gene was cloned into NotI-EcoRI-digested pMACs 4-IRES.II (contains EMCV IRES and truncated hCD4) using T4 DNA Ligase (NEB). Second, 3’ arm of homology to exon 5 was cloned into BamH1-NotI-digested PL451 (contains Neomycin cassette and loxP; NCI Frederick) using T4 DNA Ligase (NEB). Third, Step 2 was digested with EcoRI (NEB), blunted with Klenow (NEB) and then digested with ClaI (NEB). Next, Step 3 was cloned into Step 4 with T4 DNA ligase (NEB) and then this recominbeering fragment (5’ arm of homology, EMCV IRES, truncated hCD4, Neo cassette, 3’ loxP site and 3’ arm of homology) was linearized with NotI (NEB) and recombineered into RP24-227B3 BAC clone that was transformed into SW102 strain by electrophoration (186 ohms, 1.75 kV, 25 μF). Lastly, 5’ loxP site was added upstream of exon 1 using a galK cassette (NCI Frederick) that was PCR amplified with LongAmp Taq DNA polymerase (NEB), followed by recombineering-based gap repair using 5’ and 3’ arms of homology and cloning into PL253 (NCI Frederick) to generate an ES targeting construct. The *Il22*^hCD4^ targeting construct (100 ng) was linearized with NotI (NEB) and electroporated into Bruce4 mouse ES cells. Drug-resistant clones that were properly targeted for the *Il22* gene and not the pseudogene were selected (based on Southern-blot analysis; see Figure S1B) and microinjected into albino C57BL/6 blastocysts at the UAB Transgenic and Genetically Engineered Models (TGEM) Core. Founder lines were established from chimeric mice, crossed to *Actb-Flpe* mice to remove the Neomycin cassette and then bred to homozygousity. See Oligonucleotide section in Key Resources Table for detailed primer information.

### Southern Blot

Genomic DNA (gDNA) from ES clones (grown on 96-well plates) was prepared by first placing washed and aspirated plates at −80**°**C for 3 hours, followed by incubation in Lysis buffer (10 mM Tris (pH 7.5), 10 mM EDTA (pH 8.0), 10 mM NaCl, 0.5% Sarcosyl, 1 mg/ml Proteinase K) at 60**°**C overnight. The next day, 100 ul of cold Precipitation Solution (75 mM NaCl in 200 proof ethanol) was added per well and stored at RT for 1 hr to adhere DNA to the plate. Plates were then washed gently with 70% ethanol and air dried at RT for 1 hr. Plates were stored in a humidified chamber at 4**°**C. To screen the 5’ end of the targeting cassette, gDNA was digested with EcoRV (NEB) and for the 3’ end, gDNA was digested with BstXI (NEB) at 37°C for 3 hrs. gDNA was then separated on a 1% agarose gel in TBE buffer overnight at 35-40V. The gDNA/gel was then depurinated with 0.125M Hydrochloric acid (Fisher) for 10 min with gentle agitation, rinsed in deionized water, denatured in Denaturation buffer [0.5N NaOH and 1.5M NaCl (Fisher)] for 30 min with gentle agitation and then transferred to Hybond-XL membranes (GE Healthcare Amersham/Fisher). Synthesized DNA specific to regions of homologous recombination (see Oligonuleotide section in Key Resources Table) was labeled with ^32^P (Amersham) using the High Prime labeling kit (Roche) according to manufacturer’s instructions and unincorporated ^32^P was removed with G-50 Sephadex Quick Spin columns (Fisher). Prehybridization (30 min to 1 hr) followed by hybridization (overnight) were carried out at 65**°**C in Church buffer (25 ml 1M sodium phosphate buffer (pH 7.2), 17.5 ml 20% SDS, 0.1 ml 0.5M EDTA (pH 8.0), 0.5 ml salmon sperm DNA (10 mg/ml), 6.9 ml H_2_0). Probes (5 x 10^7^ cpm) were boiled for 5 min prior to hydrization. Next day, blots were washed in Washing Solution (40 ml 1M sodium phosphate buffer (pH 7.2), 50 ml 20% SDS, 910 ml H_2_0) at 60**°**C for 25 min, dried and placed in saran wrap in a phosphor cassette with an intensifying screen. The cassette was stored at −80**°**C for 1-3 days and then visualized with a Phosphorimager (Amersham). See details in Key Resources Table.

### Isolation of Intestinal Cells and Bacteria

Intestinal tissues were flushed, opened longitudinally and then cut into strips of 1 cm length. Tissue pieces were incubated for 20 min at 37**°**C with 1mM DTT (Sigma), followed by 2 mM EDTA (Invitrogen) in H5H media (1x HBSS, 5% FBS, 20 mM Hepes, and 2.5 mM 2-**β**-ME). For analysis of *C.r*-GFP released from intestinal epithelial cells (IECs), supernatant from DTT/EDTA prep were first spun at 500 rpm for 5 min at 4**°**C to remove cell debris and then spun at 8000 rpm for 15 min at 4**°**C. GFP^+^ bacteria were enumerated by flow cytometry in log scale using PKH26 reference beads (Sigma). In separate experiments and for analysis of IELs and IECs, cells from the DTT/EDTA prep were spun down at 1500 rpm for 10 minutes at 4**°**C. For isolation of lamina propria (LP) cells, tissue pieces remaining after the DTT/EDTA step were chopped and incubated for 40 min at 37**°**C with Collagenase D (2 mg/ml; Sigma) and DNase (1 mg/ml; Sigma) in R10 media (1x RPMI 1640, 10% FBS, 1x Pen/Strep, 1x NEAA, 1mM, Sodium pyruvate and 2.5 mM 2-**β**-ME). IECs and LP cells were then purified on a 40%/75% Percoll gradient by centrifugation for 20 min at 25**°**C and 600*g* with no brake. See details in Key Resources Table.

### Flow Cytometry and Cell Sorting

Colon cells were stained with Fc Block (Clone 2.4G2) followed by staining with fluorescent-labeled antibodies in FACS buffer (1x PBS and 2% FBS) on ice in 96 well round bottom plates. IECs were stained and sorted in 1x PBS with 5% FBS and 2mM EDTA to reduce cell clumping on ice in 1.5 ml microcentrifuge tubes. For intracellular staining, cells were fixed and permeabilized using BD Cytofix/Cytoperm kit (BD Bioscience). Samples were acquired on an LSRII flow cytometer (BD Biosciences) or Attune NxT flow cytometer (Life Technologies) and analyzed with FlowJo software. Cells were sorted on a BD FACS Aria (BD Biosciences). The following antibodies/reagents were used: anti-CD45 (30-F11), anti-CD45.2 (104), anti-Ceacam1/CD66a (CC1), EpCAM/CD326 (G8.8), anti-human CD4 (RPA-T4), anti-mouse CD4 (RM4-5), anti-IFN*γ* (XMG1.2), anti-IL-7R/CD127 (A7R34), anti-IL-17A (TC11-18H10), anti-IL-22 (1H8PWSR), anti-IL-33R/ST2 (RMST2-2), anti-MHC CII (I-A/I-E; 114.15.2), anti-NKp46/CD335 (29A1.4), anti-TCR*β* (H57-597), anti-TCR*γδ* (eBioGL3), Live/Dead Fixable Near-IR dead cell dye and TO-PRO-3. See details in Key Resources Table.

### Tissue Preparation

For immunostaining, LI tissues were either fixed in 2% PFA for 2 hrs at RT or in 4% PFA overnight at 4**°**C (for pSTAT3 stain). Tissue was then briefly rinsed in cold 1x PBS or put through several cold 1x PBS washes including an overnight incubation (for pSTAT3 stain), and then embedded in O.C.T. (Tissue-Tek) and frozen with 2-methyl butane chilled with liquid nitrogen. For pSTAT3 staining, tissue sections were permeabilized in cold methanol (Fisher) for 10-15 min at −20**°**C. Tissue sections were blocked at RT for 30 minutes with 10% mouse serum in 1x PBS and 0.05% Tween-20. Antibodies were diluted in 2% BSA/PBS/Tween-20 and incubated for 20-30 min at RT or ON’ at 4**°**C (pSTAT3 stain). For histological analysis, colons were segmented (proximal, middle, distal), cut longitudinally and immediately placed in 10% buffered formalin (Fisher) and processed for paraffin embedding and H&E staining. Histopathology scoring was performed in a double-blinded fashion according to published guidelines (Bleich et al., 2004). For mucin staining, colon tissue was placed in Carnoy’s fixative (60 ml ethanol, 10 ml glacial acetic acid, 30 ml chloroform) for 1 hr, followed by 95% ethanol for 1 hr and then 70% ethanol until embedded in paraffin. Antigen retrieval was performed for 15 min in the autoclave (250**°**F) using 1x Target antigen retrieval solution (Dako; S169984-2). All antibody steps were performed as described above. The following antibodies/reagents were used: anti-CD3 (17A2), anti-Ceacam1/CD66a (CC1), anti-EpCAM/CD326 (G8.8), anti-Muc1 (MH1 (CT2)), anti-Muc2 (H-300), anti-armenian hamster IgG FITC, anti-Fluorescein Alexa Fluor 488, anti-GFP Alexa Fluor 488, anti-rabbit Alexa Fluor 488, anti-rabbit Alexa Fluor 594, anti-rat Biotin, Streptavidin-Alexa Fluor 594, Prolong Gold antifade mountant with or without DAPI and UEA-1 lectin. See details in Key Resources Table.

### Microscopy

Confocal images were obtained using a Nikon A1 confocal microscope and Nikon NIS-Elements AR 4.20 program at the HRIF Imaging core at UAB. Epifluorescent and H&E images were taken using a Nikon Eclipse E800 microscope and either a Nikon Cool-SNAP Myo camera or SPOT camera, respectively. Cells and area measurements were enumerated using the Nikon NIS-Elements BR 4.5 software. LCM images were obtained using a DFC450C RGB CCD camera and Laser Microdissection LMD6 microscope (Leica Microsystems).

### Laser Microdissection

Unfixed colon tissue was frozen in O.C.T. compound (Tissue-Tek) in liquid nitrogen-cooled 2-methyl butane (Sigma). Ten micron sections were melted onto PEN-membrane glass slides (Leica) and stained with Cresyl Violet dye (Ambion). Stained epithelial cells (150,000-250,000 μm^2^ per cap) were captured into microcentrifuge tubes using a Leica LMD6 instrument. RNA was extracted using the miRNeasy Micro RNA isolation kit (Qiagen). See details in Key Resources Table.

### Real Time PCR

cDNA synthesis was performed with iScript reverse transcription (RT) Supermix (Bio-Rad) according to manufacturer’s instructions. cDNA amplification was analyzed with SsoAdvanced Universal SYBR Green Supermix (Bio-Rad) in a Biorad CFX qPCR instrument. The following primers (generated by IDT) were used: Muc2, AAGTGGCATTGTGTGCCAACCA (forward) and TGCAGCACTTGTCATCTGGGTT (reverse); Scnn1a, TGGGCAGCTTCATCTTTAC (forward) and CCAGAGATTGGAGTTGTTCTT (reverse); Slc12a2, CATACACTGCCGAGAGTAAAG (forward) and CCACGATCCATGACAATCTAA (reverse). See details in Key Resources Table.

### Fecal pellet Extracts

Fecal pellets were collected in 1x PBS with Complete protease inhibitor cocktail (Roche), vortexed and spun at 13,000 rpm for 10 minutes at 4**°**C to remove debris. Clarified supernatant was transferred to a clean microfuge tube and stored at −20**°**C. See details in Key Resources Table.

### IgG ELISAs

A 96-well high-binding assay plate (Corning) was coated with 10 Dg/ml of unlabeled polyclonal anti-mouse Ig (H+L) in 1x PBS overnight at 4**°**C. After 4 washes with 1x PBS, 1% BSA in 1x PBS Blocking solution was added to the plate and incubated at RT for 1 hr. After 4 washes with 1x PBS/Tween-20, samples were diluted in 1% BSA/PBS/Tween- 20 and incubated at RT for 2 hrs. After 4 washes with 1x PBS/Tween-20, 1:4000 dilution of anti-mouse IgG (H+L)-HRP conjugated was incubated at RT for 2 hrs. After the final 4 washes, 100 Dl of a TMB single solution (Life Tech) was added to the plate and the chemiluminescence signal was stopped with 2N Sulfuric acid and read at 450 nm. All samples were run in triplicate and IgG standards were included on all plates. The following antibodies purchased from Southern Biotech were used: anti-mouse Ig (H+L) unconjugated (for coating plate), purified mouse IgG (for standard) and anti-mouse IgG HRP labeled (for detection). See details in Key Resources Table.

### RNA-sequencing and Data Analysis

Three subsets of colonic epithelial cells were isolated from naïve and D9 *C.r*-infected *Il22*^hCD4^ (Control) and *Il22*^ΔTcell^ (T cell cKO) mice. Tissue was pooled from 2-3 mice per condition and cells were sorted directly into Trizol LS. Total RNA was extracted using Trizol LS/Chloroform extraction (Invitrogen/ThermoFisher) and miRNeasy Micro2.5. Kit (Qiagen) according to the manufacturer’s protocol. Next, 30 ng of total RNA was used to prepare mRNA-seq library using the Nugen Universal Plus mRNA-seq library prep kit (Nugen), and libraries were sequenced on the SE50 Illumina 2500 in Rapid Run Mode. Single-end reads with lengths of 50 nucleotides (∼20M reads per condition) were generated for subsequent bioinformatics analysis. Adaptors were trimmed and aberrant reads were removed using TrimGalore (version 0.4.5). The quality controlled reads were mapped onto the mouse genome build GRCm38 (ENSEMBL.mus_musculus.release-75) (Hubbard et al., 2002) using STAR (version 2.5.3) (Dobin et al., 2013). BAM files were sorted using SAMtools (version 0.1.18) (Li et al., 2009), and reads were counted for each gene using HTSeq (version 0.7.2) (Anders et al., 2015). Differential expression analysis was performed using DESeq2 (version 1.18.1) (Love et al., 2014) using R (version 3.4.3). Dispersion shrinkage of fold changes was performed with the ASHR algorithm (Stephens, 2017). The fgsea R package (version 1.4.0) (Sergushichev, 2016) was used for gene set enrichment. Gene sets used include those from the mouse cell atlas of the small intestine (Haber et al., 2017a) and MSigDB (Liberzon et al., 2015; 2011; Subramanian et al., 2005). Data is publicly available in GEO (Barrett et al., 2013) under GSE114338.

## Extended Bioinformatics Methods

### Pre-processing of RNA-sequencing data

The pre-processing pipeline was operated through the UNIX shell. Each raw fastq file was processed using the default settings of each analysis tool except as specified below. Standard Illumina adaptors were used to trim reads with TrimGalore. The ENCODE options for standard long RNA-seq were utilized for mapping using STAR, as defined in the STAR manual (Dobin et al., 2013). Quality of libraries was assessed by overall mapping rate, and libraries with less than 70% mapping rate were discarded from further analysis.

TrimGalore (version 0.4.5) *--illumina*

STAR (version2.5.3) *--outSAMtype BAM SortedByCoordinate Unsorted --outFilterType BySJout --outFilterMultimapNmax 20 --alignSJoverhangMin 8 –alignSJDBoverhangMin 1 --outFilterMismatchNmax 999 --outFilterMismatchNoverReadLmax 0.04 -- alignIntronMin 20 --alignIntronMax 100000 --alignMatesGapMax 100000* HTSeq (version 0.7.2) *--stranded=yes -t exon*

## Statistical analysis of RNA-sequencing data

The R software environment (version 3.4.3) with BioConductor (version 3.6) was used for statistical analysis. The random seed was set to 15144305.

### Exploratory analysis and Quality Control

Gene counts were filtered for genes catalogued by ENSEMBL (ENSEMBL.mus_musculus.release-75) as *protein_coding*. We subjected cleaned sequencing data to a battery of exploratory analyses using the R package DESeq2 (version 1.18.1) (Love et al., 2014) in order to inform model selection for analysis of differential expression, including distance matrices and principal component plots of the top 500 differentially expressed genes after rlog transformation (Love et al., 2014). We visually inspected these together in order to identify low-quality samples, outliers, and unexpected trends relating to batch ID or other important covariates to be used in modeling. Additional QC measures were carried out as described in the DESeq2 vignette (http://bioconductor.org/packages/devel/bioc/vignettes/DESeq2/inst/doc/DESeq2.html).

### Count Plots

Counts were normalized using the sequencing depth and plotted on the y-axis on a log_10_- scale (**Figure 6E**). In each count plot, different groups are plotted along the x-axis, while these normalized counts are plotted on the y-axis (log_10_-scale). The means of these groups were used to generate the endpoints for each line plotted; as such the lines visualize the mean difference in normalized expression between pairs of groups.

### Differential Gene Expression Analysis

We tested for differential expression using a negative binomial generalized linear model as described previously (Love et al., 2014). We modeled the effect of IL-22-deficient T cells during infection in each location tested (small crypt cell samples, large crypt cell samples, surface cell samples) separately (1) and then pooled naive cells from all of these locations and examined the effect of IL-22-deficient T cells in pooled naive and infected intestinal epithelial cells (2). We also modeled the effect of infection itself (i.e., of naive versus infected cells) in small crypt samples (3), large crypt samples (4), surface samples (5), and pooled control samples (6) and pooled samples from D9 *C.r*-infected *Il22*^ΔTcell^ (T cell cKO) (7). Next, we tested the effect of location on gene expression within the intestinal crypt (surface, small crypt, or large crypt) in naive (8) and infected cells (9). Finally, we were interested in whether cells from D9 *C.r*-infected Control and *Il22*^ΔTcell^ mice demonstrate cell-type specific differences.

Generally, Wald Tests were selected in place of likelihood ratio tests and stratified analysis outperformed pooled analysis for most models. Final model specifications for each of these tests are available by reasonable request. We set alpha, our allowable type I error rate, at a 0.05 after correction for multiple testing using independent hypothesis weighting (Ignatiadis et al., 2016). For each model, we assessed model fit and model validity based on dispersion estimates, overall type I error rate, Cooks’ distances, inspection of count-dispersion plots and MA plots (log_2_ fold change versus log_2_ (base mean/normalized read counts) for competing models for each research question.

### MA plots

MA plots were generated using ggplot2 (Wickham, 2009) (**Figures S5E-G**). Shrunken log_2_ fold change (y-axis) in gene expression attributable to the variable of interest were plotted against the mean of normalized counts for all the samples to generate MA plots. Ashr were used as shrinkage estimators in order to aid visualization of log fold change data (Love et al., 2014; Stephens, 2017). This was in turn used for the assessment of model validity and performance. For these plots, as elsewhere, coloration (in red) indicates that the adjusted p-value (from the linear modeling step, above) of an observation was less than alpha, set at an FDR of 0.05. Color saturation was also based on adjusted *p*-value; observations with *p_adj_* < 0.05 were fully saturated, with saturation decreasing to 0.2 as *p_adj_* increases to 1. Genes were labeled using the ggrepel package in R.

### Gene Set Enrichment Analysis (GSEA)

To prepare data for gene set enrichment analysis, results of differential gene expression analysis were ranked by signed *p*-value. Gene sets from the small intestine mouse cell atlas (Haber et al., 2017b) were curated by taking the union of signature marker genes for each cell type from droplet and plate-based datasets (Custom Pathways). Interferon and tumor necrosis factor gene sets were obtained from the Hallmark Gene Set Collection (Liberzon et al., 2015) and from the ImmGen interferon transcriptional network (Mostafavi et al., 2016). Additionally, three pathways that represent immune defense genes (GO_cellular_defense_response, GO_innate_immune_response, and GO_regulation_of_defense_response) were selected from the Gene Ontologies collection (Ashburner et al., 2000; The Gene Ontology Consortium, 2017). Publicly available datasets from experiments utilizing IL-22 in intestinal tissue were used to create custom IL-22-related gene sets. When possible, analyzed data from original publications were used to create gene sets; however, when this data was not available, raw transcriptomic data was analyzed in a manner consistent with the reported methods. The limma R package (Ritchie et al., 2015) was used to analyze data from GSE15955 (Pickert et al., 2009) and GSE10010 (Zheng et al., 2008) for differential expression. For GSE15955, up- and down-regulated genes were selected that had a fold-change greater than 2 and adjusted p-value less than 0.1. For GSE10010, the top 250 genes by adjusted *p*-value were selected and divided into up-regulated and down-regulated gene sets. Additionally, IL-22-regulated genes identified in *Il22ra1*^−/−^ mice (Pham et al., 2014) were obtained from supplemental data to create a gene set of IL-22-induced genes.

A *p*-value quantifying the likelihood that a given gene set displays the observed level of enrichment for DE genes was calculated using Fast Gene Set Enrichment Analysis (*fgsea*, version 1.4.0) (Sergushichev, 2016) with 1 million permutations. Gene set enrichment p-values of Normalized Enrichment Scores (NES) were corrected with the Benjamini-Hochberg procedure (Benjamini and Hochberg, 1995).

### Barcode Plots, Dot Plots, and 2-way Scatterplots

We used the *plotEnrichment* (Sergushichev, 2016) function from the *fgsea* package to generate barcode plots (**Figure 6H**). The Random Walk curves were recolored in Adobe Illustrator according to sign of NES value. We generated dot plots of NES values using *ggplot2* (Wickham, 2009) (**Figures 6F-G**). Coloration of points was based on a continuous gradient from most negative to most positive NES, with the transition from blue to red coloration at NES = 0. The size of dots is directly proportional to −log_10_ of the adjusted p-value generated from the enrichment of each pathway for each comparison. 2-way plots were made by plotting DEGs (adjusted p-value < 0.01) on the x-axis (red), y-axis (blue) or significant in x-axis and y-axis (purple) (**Figure 6D**).

### Pathway Enrichment analysis and Pathway-to-Pathway m-type network construction

Wikipathways, KEGG pathways and Custom pathways (see GSEA section above) were used for identifying DEGs and Pathways in D9 *C.r*-infected *Il22*^ΔTcell^ (T cell cKO) vs D9 *C.r*-infected *Il22*^hCD4^ (Control) mice (**Figure 7** and **Figure S6**). First, we performed an enrichment analysis using a hypergeometric test as implemented in PAGER 2.0 (Yue et al., 2018). The false discovery rate (FDR) was set to 0.05 using Benjamini–Hochberg procedure (Benjamini and Hochberg, 1995). Second, redundant pathways were identified using two metrics: pathway membership similarity and content similarity. Specifically, all pairs of pathways were tested for similarity of pathway membership using the Jaccard index, and the cutoff was set to 0.7. In addition, content similarity was assessed using the Jaccard index of pathway names’ bigrams, and the cutoff was again set to 0.7, retaining the pathway that contained a larger proportion of the genes. Following this, we established m-type pathway-to-pathway relationships using the hypergeometric test in PAGER 2.0 and set the FDR cutoff to 0.01, then these networks were visualized using Cytoscape software (Shannon et al., 2003). In order to annotate the m-type networks, we conducted an enrichment analysis. Interactors of IL-22 were identified by querying Protein-Protein Interactions in the STRING database (Szklarczyk et al., 2017) with a score ≥0.85. Then, we performed enrichment analysis using IL-22 and its interactors and set alpha at 0.05. By taking the union of enriched pathways from infected (*C.r* D9) and naive samples, we identified a network of pathways relevant to the infection. Finally, we evaluated the importance of each pathway by computing a final score (fs), which was set equal to the distance-based sum of all pairwise samples log_2_ fold change (aggregated at the pathway level) alongside the distance-based sum of the pairwise samples’ Enrichment Scores (ES).

## Quantification and Statistical Analysis

All statistical tests were done with GraphPad Prism software. The appropriate statistical test for each experiment is noted in the figures. For all graphs, bars or lines mean and error bars indicate s.e.m. For data plotted on log scale, log-transformed data was compared. Statistical significance was calculated with either ANOVA or the nonparametric Mann-Whitney test. One-way ANOVAs were run with post-hoc Tukey tests. Two-way ANOVAs were run with Bonferroni posttests.

### Data and Code Availability

Raw and process data files for RNA sequencing analysis have been deposited in the NCBI Gene Expression Omnibus under accession number GEO: GSE114338.

## ACKNOWLEDGMENTS

We gratefully acknowledge Dr. T.R. Schoeb and the Comparative Pathology Laboratory at UAB for histology scoring and tissue processing, respectively, and J. Day (La Jolla Insitute for Immunology) for assistance with. We thank Dr. R.A. Kesterson and the UAB Transgenic and Genetically Engineered Models (TGEM) Core for ES cell microinjection, and Marion Spell and the UAB CFAR Flow core for the sorting of IECs. We also recognize Shawn Williams and the HRIF Imaging core at UAB for assistance with confocal microscopy. We also thank the Laser Capture Microdissection core and the Small Animal Imaging Shared Facility at UAB. This work was supported by grants from the NIH and Crohn’s and Colitis Foundation (C.T.W.), T32 training grant funds from NIH/NIAID and the UAB Training Program in Immunologic Diseases and Basic Immunology (C.L.Z., D.P., D.J.S.), UAB MSTP training grant funds (J.R.S) and UAB NIH F30 grants (J.R.S, C.E.M.)

## Author Contributions

C.L.Z. and C.T.W. conceptualized the project, designed the experiments, interpreted the results and wrote the manuscript. C.L.Z. performed most of the experiments. C.L.Z. and H.T. generated the *Il22^hCD4^* reporter/cKO mouse line. D.J.S. and J.R.S. assisted with the bioluminescence imaging experiments and B.C. performed the LCM experiments. S.H., D.J.S., J.R.S. and C.E.M. provided discussion and advice on experiments. R.T.H. assisted with RNA-seq experiments. S.J.W., V.A.L., M.G. and D.P. carried out bioinformatics analyses and assisted with RNA-seq data visualization. Y.Z. carried out pathway analysis.

## Declaration of Interests

The authors declare no competing interests.

## Notes

### Competing Interest Statement

The authors have declared no competing interest.

