## Supplemental Materials for "A Non-redundant Role for T cell-derived IL-22 in Antibacterial Defense of Colonic Crypts"

**Supplemental Information**

**Figure S1. *Il22*$^{hCD4}$ construct design and characterization. Related to Figure 1.**
(A) *Il22*$^{hCD4}$ mice were generated using a targeting cassette containing an EMCV IRES, truncated human CD4 (hCD4) gene, and a frt-flanked (◊) neomycin resistance cassette. LoxP (O) sites were placed upstream of exon 1 and into the fifth exon of the *Il22* gene, immediately 3' of the stop codon (*) and 5' of the 3' untranslated region of exon 5. The targeting cassette was transfected into Bruce4 mouse ES cells and Ganciclovir/G418-resistant clones were screened for correct targeting of the *Il22* gene by Southern blot analysis (B) (N, non-targeted; A, *Il22* allele targeted; B, *Iltifb* allele targeted). (C) T cells from WT or *Il22*$^{hCD4}$ mice were differentiated *in vitro* with IL-6 (10 ng/ml) and IL-23 (10 ng/ml) for 4-5 days. T cells were restimulated with PMA/Ionomycin and BrefeldinA, and then surface stained for TCRβ, mCD4, hCD4, and Live/Dead dye, followed by intracellular staining for IL-22 and analyzed by flow cytometry; mCD4$^{-}$ cells (red), mCD4$^{+}$ cells (blue). (D) Colon cells from *C.r*-infected *Il22*$^{hCD4}$ mice were stimulated with PMA/Ion and IL-23 for 4 hrs and then stained for TCRβ, hCD4, mCD4, TCRγδ and L/D dye and analyzed by flow cytometry. (E-F) IELs and LPLs from small intestine, cecum and colon of D9 *C.r*-infected *Rorc*$^{EGFP}$ *Il22*$^{hCD4}$ mice were stimulated with rmIL-23 for 4 hrs and then stained for hCD4, IL-7R, IL-33R, and Live/Dead dye and analyzed by flow cytometry with total cells (E) gated on live IL-7R$^{+}$ IL-33R$^{-}$ cells and (F) gated on live IL-7R$^{+}$ IL-33R$^{-}$ RORγ$^{+}$ cells. Numbers in (F) represent percentages of hCD4 (IL-22$^{+}$) innate cells (TCRβ$^{-}$) or T cells (TCRβ$^{+}$). (G) IELs and LPLs from small intestine, cecum and colon of D9 *C.r*-infected *Rorc*$^{EGFP}$*Il22*$^{hCD4}$ mice were stimulated with rmIL-23 for 4 hrs and then stained with for hCD4, IL-7R, IL-33R, NKp46 and L/D dye and analyzed by flow cytometry (gated on live IL-7R$^{+}$ IL-33R$^{-}$ cells). Numbers represent percentages of hCD4 (IL-22$^{+}$) NK cells (NKp46$^{+}$). Data shown are concatenated plots from 3 mice.
3-4 mice per time point, 3 independent experiments.

**Figure S2. Colonization kinetics of *C.r* in the large intestine and Characterization of *Il22*$^{Plzf}$ Innate cell cKO mice. Related to Figure 2.**
(A) *C.r* from supernatants of IEC preps of the distal SI (ileum) and LI (cecum, proximal colon, middle colon, and distal colon) from naïve and D1-D3 *C.r*-GFP-infected BL/6 mice was stained with TO-PRO-3 and analyzed by flow cytometry in log scale. (B) *C.r* from supernatant of IEC preps of cecum and total colon from naïve and *C.r*-infected BL/6 mice at various time points were stained with TO-PRO-3 and analyzed by flow cytometry in log scale. (C) GFP$^{+}$ TO-PRO-3$^{+}$ *C.r* was quantitated by flow cytometry using PKH26 reference beads (Sigma). Two-way ANOVA; $**p < 0.01$ and $***p < 0.001$

comparing naïve and infected. nd=not detected. 3 mice per time point, 2 independent experiments. Consistent with pioneering studies (Wiles et al., 2004), *C.r* colonizes the cecum within 24 hrs, followed by colonization in the colon within 3-4 days after inoculation. In addition, *C.r* is cleared from the cecum prior to the colon. Therefore, flow cytometric quantitation allows an accurate assessment of the number of *C.r* attached to epithelial cells from different regions of the colon. (D) Colon LP cells from *C.r*-infected *Il22*$^{hCD4}$ (Cntrl) and *Il22*$^{Plzf}$ (Innate cell cKO; green) mice were stimulated with rmIL-23 for 4 hrs and then stained for hCD4, TCRβ, mCD4, TCRγδ□and Live/Dead dye and analyzed by flow cytometry. Two-way ANOVA; **p* <0.05, ***p* <0.01 comparing Cntrl and cKO mice. 2-3 mice per group, 2 independent experiments. (E) LI tissues from d8 *C.r*-infected *Il22*$^{hCD4}$ and *Il22*$^{Plzf}$ (Innate cell cKO; green) mice were stained with hematoxylin and eosin for blinded histology scoring. ns=not significant. 3-5 mice per group, 2 independent experiments.

**Figure S3. *Il22*$^{EIIa}$ global KO mice have enhanced crypt loss at the peak of *C.r* infection. Related to Figure 3.**

(A) LI tissues (cecum, proximal colon, middle colon, and distal colon) from d8 *C.r*-infected *Il22*$^{hCD4}$ (WT) and *Il22*$^{EIIa}$ (global KO; red) mice was stained with hematoxylin and eosin for blinded histological scoring of epithelial cell hyperplasia, goblet cell loss and epithelial cell degeneration/crypt loss. (B) LI tissues from d8 *C.r*-infected *Il22*$^{hCD4}$ and *Il22*$^{Plzf}$ (Innate cell cKO; green) mice were stained with hematoxylin and eosin for blinded histological scoring of epithelial cell hyperplasia, goblet cell loss and epithelial cell degeneration/crypt loss. Two-way ANOVA; **p* <0.05, ***p* <0.01, ****p* <0.001 comparing Cntrl and cKO mice. 3-5 mice per group, 2 independent experiments.

**Figure S4. *Il22*$^{\Delta Tcell}$ T cell cKO mice have exacerbated pathology and decreased *Muc1* expression on crypt IECs during late phase of *C.r* infection. Related to Figure 4.**

(A-B) LI tissues (cecum, proximal colon, middle colon, and distal colon) from d16 *C.r*-infected *Il22*$^{hCD4}$ (Cntrl) and *Il22*$^{\Delta Tcell}$ (T cell cKO) mice was stained with hematoxylin and eosin for blinded histological scoring of (A) total histology scoring middle and distal colon and (B) epithelial cell hyperplasia, goblet cell loss and epithelial cell degeneration/crypt loss. *Il22*$^{\Delta Tcell}$ mice have prolonged hyperplasia and crypt loss during the last phase of *C.r* infection when control mice have mostly cleared *C.r* from superficial IECs. Two-way ANOVA; **p* <0.05 and ****p* <0.001 comparing Cntrl and

*Il22*$^{\Delta Tcell}$ mice. 3-5 mice per group, 2 independent experiments. (C) Supernatant from fecal extracts was analyzed for total IgG levels by ELISA. 3-4 mice per group, 2 independent experiments. t-test; $^{**}p$ <0.01 comparing Cntrl and *Il22*$^{\Delta Tcell}$ mice. ns=not significant. Consistent with elevated serum IgG levels in *Il22ra1*$^{-/-}$ mice (Pham et al., 2014), fecal IgG levels were elevated in *Il22*$^{\Delta Tcell}$ mice indicating that IL-22$^{+}$ T cell-driven crypt protection may not depend on antibody-mediated bacterial clearance. (D) Colon tissue from *C.r* D12 *Il22*$^{hCD4}$ (Cntrl) and *Il22*$^{\Delta Tcell}$ (T cell cKO) mice was stained for Mucin1 (Muc1; green), Mucin2 (Muc2; red) or Ulex Europaeus Agglutinin I Lectin (UEA1; green) and DAPI (blue). 3-4 mice per group. In accord with previous studies (Bergstrom et al., 2008; Lindén et al., 2008), *C.r*-infected control mice exhibited increased Muc1$^{+}$ and decreased Muc2$^{+}$ expression on crypt IECs. Since Muc1 and Muc2 play protective roles in bacterial clearance in the intestines (Bergstrom et al., 2010; Lindén et al., 2009) T cell-derived IL-22-driven upregulation of Muc1 on the apical surface of colonic goblet cells likely plays a crucial role in crypt defense by hindering bacterial growth within the crypts during *C.r* infection. (E) Mid/distal colon epithelial cells from naïve and *C.r*-infected BL/6 mice were stained for EpCAM1, MHC CII, CD45 and Live/Dead dye and analyzed by flow cytometry. 4 mice per group. One-way ANOVA; $^{**}p$ <0.01 and $^{***}p$ <0.001 comparing naïve and infected groups. MHC CII has been show to be expressed by colonic IECs in a murine adoptive T cell transfer colitis model (Thelemann et al., 2014). At steady state, surface IECs isolated from colons of naïve mice express *H2-Ab1* mRNA (Figure 6E) with approximately 5% of IECs expressing MHC CII protein. Interestingly, as the T cell response increases around day 8-9 after *C.r* inoculation, *H2-Ab1* mRNA is upregulated on SC and LC IECs (Figure 6E) with 20% of colonic IECs expressing MHC CII protein. Strikingly, on day 12 when IFNγ responses are heightened, up to 60% of all colonic IECs express MHC CII. Together these data suggest that MHC II upregulation on colonic crypt IECs may contribute to antigen presentation to CD4 T cells for eradication of *C.r* and protection of the crypts.

**Figure S5. IL-22$^{+}$ T cells amplify genes involved in host defense and reduce genes induced by IFNγ. Related to Figure 6.**

(A) Cells from LI tissues (cecum, proximal colon, middle colon and distal colon) were stained for EpCAM1, CEACAM1, CD45 and Live/Dead (L/D) dye and analyzed by flow cytometry. (B) RNA-seq was performed on sorted small crypt (SC) (EpCAM1$^{+}$ CEACAM1$^{lo}$ FSC$^{lo}$ SSC$^{lo}$ CD45$^{-}$ L/D dye$^{-}$; blue), large crypt (LC) (EpCAM1$^{+}$ CEACAM1$^{int}$ FSC$^{int}$ SSC$^{int}$ CD45$^{-}$ L/D dye$^{-}$; green) and surface IECs (Srf) (EpCAM1$^{+}$ CEACAM1$^{hi}$ FSC$^{hi}$ SSC$^{hi}$ CD45$^{-}$ L/D dye$^{-}$; red) from mid/distal colon to determine relative expression of *Ceacam1*, *Il22ra1*, *Il10rb* (pairs with *Il22ra1* to form IL-22R signaling complex), *Slc12a2* (sodium/potassium/chloride transporter expressed by crypt cells (Pena-Munzenmayer, 2005)), *Muc2* (mucin gene expressed by goblet cells (Allen et al., 1998)) and *Scnn1a* (sodium channel expressed by luminal surface IECs (Duc et al., 1994) from naïve BL/6 mice. Counts were normalized by library size. **p*adj <0.1, ***p*adj <0.01, ****p*adj <0.001 comparing gene expression between IEC subsets. 2-3 mice per sample, 1-2 independent experiments per naïve group and 3-4 independent experiments per infected group. (C) Colon tissue from naïve BL/6 mice was stained with Cresyl Violet dye and 3 regions of the colon: crypt base (blue), crypt neck (green) and superficial (red) were isolated using laser capture microdissection (LCM). (D) RNA from laser-captured regions of colon tissue were extracted and analyzed by RT-PCR for *Slc12a2*, *Muc2* and *Scnn1a*. Relative mRNA levels were normalized to *Gapdh* and the numbers indicate the quotient of specific gene per *Villin* control mRNA levels. One-way ANOVA; **p* <0.05, ***p* <0.01, ****p* <0.001 comparing gene expression between different epithelial regions. Data are representative of 3-4 independent experiments. As expected, *Ceacam1* and *Scnn1a* mRNA expression is higher on Srf IECs compared to crypt IECs. In contrast, the crypt cell marker *Slc12a2* is higher on crypt IECs compared to surface IECs. In addition, Muc2$^{+}$ cells are found in both the LC and Srf IECs suggesting that goblet cell subsets may have varying levels of *Ceacam1* expression dependent upon their stage of maturation. Interestingly, *Il22ra1* and *Il10rb* mRNA levels are 2-5 fold higher in the Srf IECs compared to crypt IECs indicating that IL-22R expression is highest on mature IECs in the colon. (E-G) MA plots of fold change ($log_2$) versus normalized read counts ($log_2$) show DEGs from SC, LC and Srf IECs from (E) naïve Cntrl versus *C.r* D9 Cntrl, (F) naïve *Il22*$^{\Delta Tcell}$ versus *C.r* D9 *Il22*$^{\Delta Tcell}$ and (G) *C.r* D9 Cntrl versus *C.r* D9 *Il22*$^{\Delta Tcell}$ mice. 2-3 mice per sample, 1-2 independent experiments per naïve group and 3-4 independent experiments per infected group. Several IL-22-inducible genes (e.g., *S100a8*, *Cxcl5*, *Lrg1*, *Tac1*) are upregulated in all IECs from *C.r* D9 Cntrl compared to naïve mice and significantly reduced in infected *Il22*$^{\Delta Tcell}$ compared to infected controls. Furthermore, *Lbp*, *Lcn2* and *Tifa* are significant DEGs in crypt cells only (SC and LC) suggesting that these genes may play an important role in IL-22-mediated T cell-driven crypt protection. In contrast, infected *Il22*$^{\Delta Tcell}$ mice displayed enhanced expression of several IFNγ-inducible genes (e.g., *Gbp2*, *Igtp*, *Cxcl9*) in all IECs compared to infected controls indicating that IL-22/IL-22R/Stat3 signals act to dampen Stat1-driven pro-inflammatory responses in colonic IECs during *C.r* infection.

**Figure S6. Pathway network analysis identifies DEGs mapping to Enterocyte-specific, oxidative stress and metabolic pathway genes. Related to Figure 7.**

(A-C) RNA-seq was performed on sorted colon IECs from D9 *C.r*-infected *Il22*$^{hCD4}$ (Cntrl) and *Il22*$^{\Delta Tcell}$ (T cell cKO) mice. (A) Heatmap of the average expression of significant DEGs in Wiki (W), KEGG (K) and Custom IEC (C) pathways was generated using Pager 2.0 (Yue et al., 2015; 2018). Color represents DEGs upregulated in *C.r* D9 Cntrl mice (red) and DEGs upregulated in *C.r* D9 *Il22*$^{\Delta Tcell}$ mice (blue). Color gradient represents the average of the DEGs ($log_2$ fold change). Numbers in the graph represent the DEG hits per size of the pathway. (B) Co-membership network was generated using Pager 2.0 (Yue et al., 2018; 2015) to display the comparison between DE pathways in small crypt (SC), large crypt (LC) and surface (Srf) IECs from *C.r* D9 Cntrl and *C.r* D9 *Il22*$^{\Delta Tcell}$ mice. Color inside the symbol depicts fold change ($log_2$, pathway upregulated in Cntrl (red), pathway upregulated in *Il22*$^{\Delta Tcell}$ (blue)), size of the symbol depicts enrichment score, shape of the symbol depicts source (Custom Pathway (square), KEGG (circle), Wiki Pathway (diamond)), and border color depicts type of interaction (Infection (green), IL-22 (orange), IL-22 interactors (pink)). Numbers represent reference identification (Ref ID) of pathways. (C) Table of the Wiki (W), KEG (K) and IEC Custom (C) pathways was generated with the pathways ranked by the final score. The final score is calculated by the sum of the pairwise distance of the sample log2 Fold Change multiplied by the sum of the pairwise distance of the sample Enrichment Score. The original links to the WikiPathways can be retrieved by www.wikipathways.org/index.php/Pathway:[GS_ID], and the original links to the KEGG Pathways can be retrieved by www.genome.jp/kegg-bin/show_pathway?[GS_ID].

2-3 mice per sample, 3-4 independent experiments per group. As expected, DEGs in IL-22/Stat3 pathways (cust4, cust28, cust30, cust31) are heightened in all IEC subsets (small crypt, large crypt, surface) from *C.r* D9 Cntrl mice compared to *C.r* D9 *Il22*$^{\Delta Tcell}$ mice. In addition, DEGs associated with pancreatic secretion (mmu04972), and fat digestion and absorption (mmu04975) (e.g., *Pla2g2a, Pla2g2f* and *Pla2g5*) are enriched in large crypt and surface IECs. DEGs associated with amino acid metabolism (e.g., *Gsr*, *Mdh2*, *Tat*) are also enriched in LC IECs from *C.r* D9 Cntrl mice. In contrast, DEGs in IFN$\alpha$ and IFN$\gamma$ pathways (cust2, cust3, WP1253) are increased in *C.r* D9 *Il22*$^{\Delta Tcell}$ mice compared to controls. Moreover, in the absence of T cell-derived IL-22, DEGs associated with glutathione metabolism and oxidative stress (e.g., *Gpx3*, *Gclc*, *Hmox1*) is heightened in small crypt and large crypt IECs. This is likely due to bacterial overgrowth in the crypts and consequently may contribute to the enhanced pathology observed in colons of *Il22*$^{\Delta Tcell}$ mice (**Figures S4A** and **S4B**).

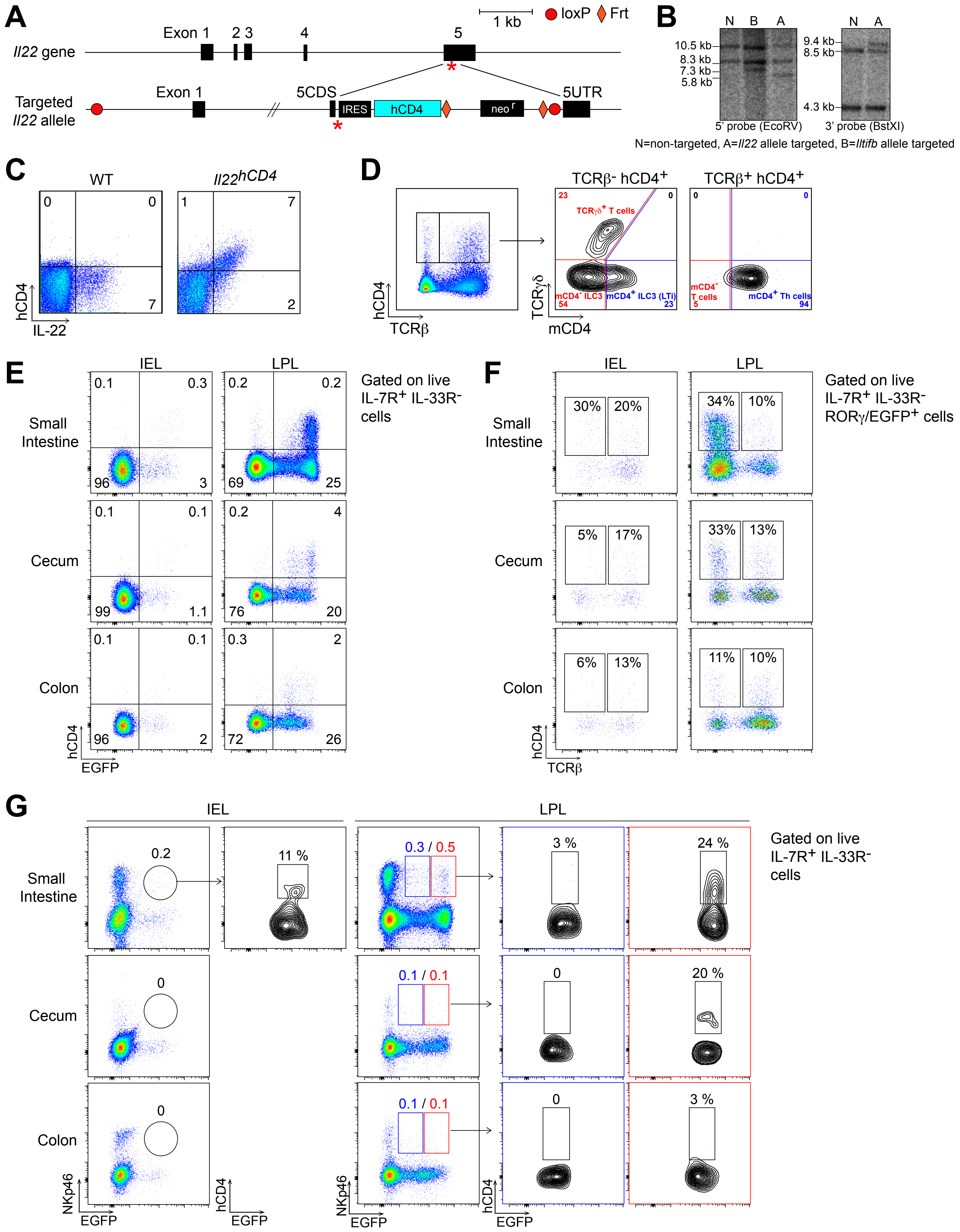

A
1 kb
loxP
Frt
Exon 1
2 3
4
5
Il22 gene
Targeted Il22 allele
Exon 1
5CDS
IRES
hCD4
neo r
5UTR
B
N B A
10.5 kb
8.3 kb
7.3 kb
5.8 kb
N A
9.4 kb
8.5 kb
4.3 kb
5' probe (EcoRV)
3' probe (BstXI)
N=non-targeted, A=Il22 allele targeted, B=Iltifb allele targeted
C
WT
Il22hCD4
hCD4
IL-22
D
TCRβ- hCD4+
TCRβ+ hCD4+
TCRγδ+ T cells
mCD4- ILC3
mCD4+ ILC3 (LTi)
mCD4- T cells
mCD4+ Th cells
hCD4
TCRβ
TCRγδ
mCD4
E
IEL
LPL
Small Intestine
Cecum
Colon
Gated on live IL-7R+ IL-33R- cells
hCD4
EGFP
F
IEL
LPL
Small Intestine
Cecum
Colon
Gated on live IL-7R+ IL-33R- RORγ/EGFP+ cells
hCD4
TCRβ
G
IEL
LPL
Small Intestine
Cecum
Colon
Gated on live IL-7R+ IL-33R- cells
NKp46
EGFP
hCD4
EGFP
NKp46
EGFP
hCD4
EGFP


**Figure S1**

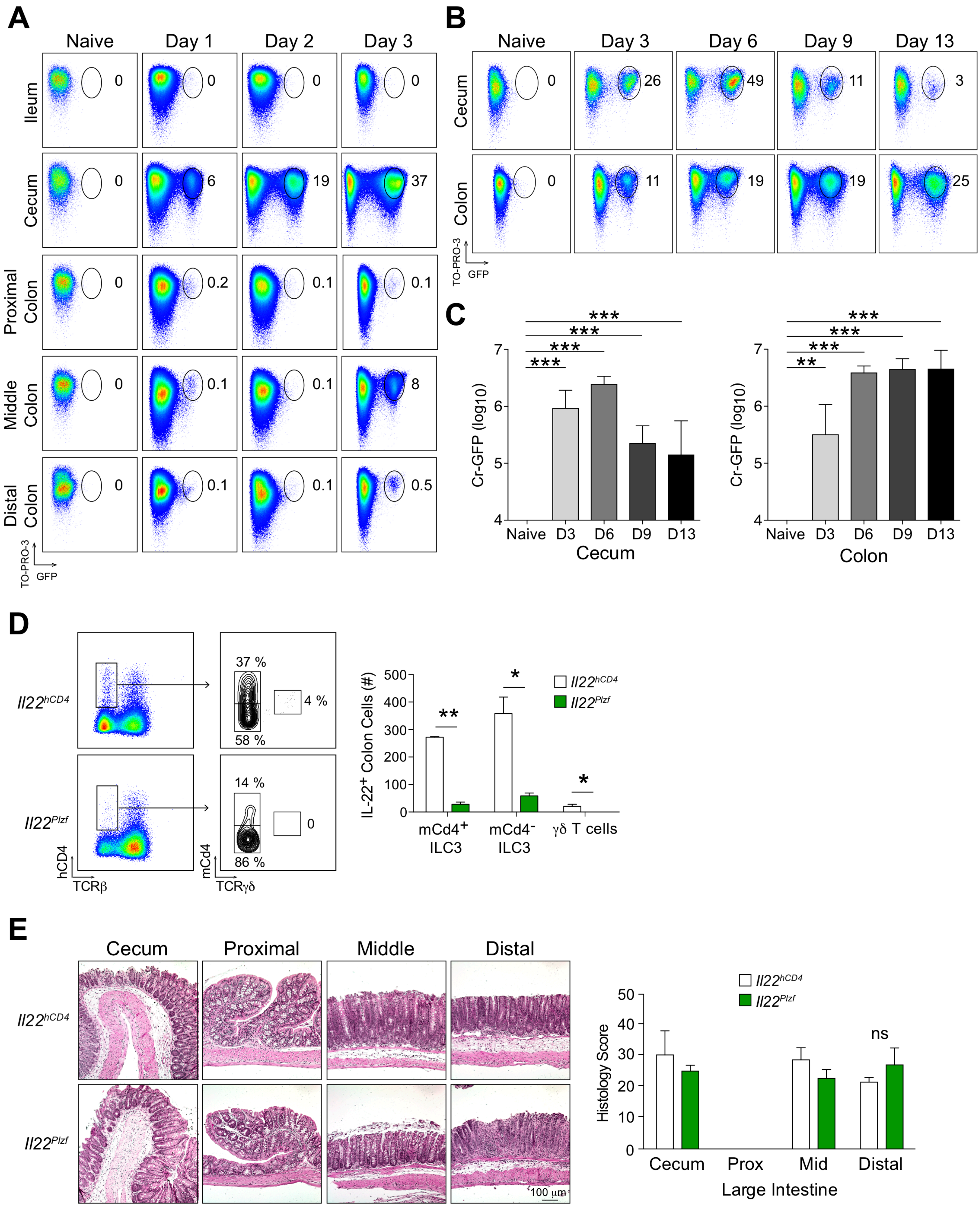

A
Naive
Day 1
Day 2
Day 3
Ileum
Cecum
Proximal Colon
Middle Colon
Distal Colon
TO-PRO-3
GFP
B
Naive
Day 3
Day 6
Day 9
Day 13
Cecum
Colon
C
Cr-GFP (log10)
Naive
D3
D6
D9
D13
Cecum
Colon
D
Il22hCD4
Il22Plzf
hCD4
TCRβ
mCd4
TCRγδ
IL-22+ Colon Cells (#)
mCd4+ ILC3
mCd4- ILC3
γδ T cells
E
Cecum
Proximal
Middle
Distal
100 µm
Histology Score
Cecum
Prox
Mid
Distal
Large Intestine
ns


**Figure S2**

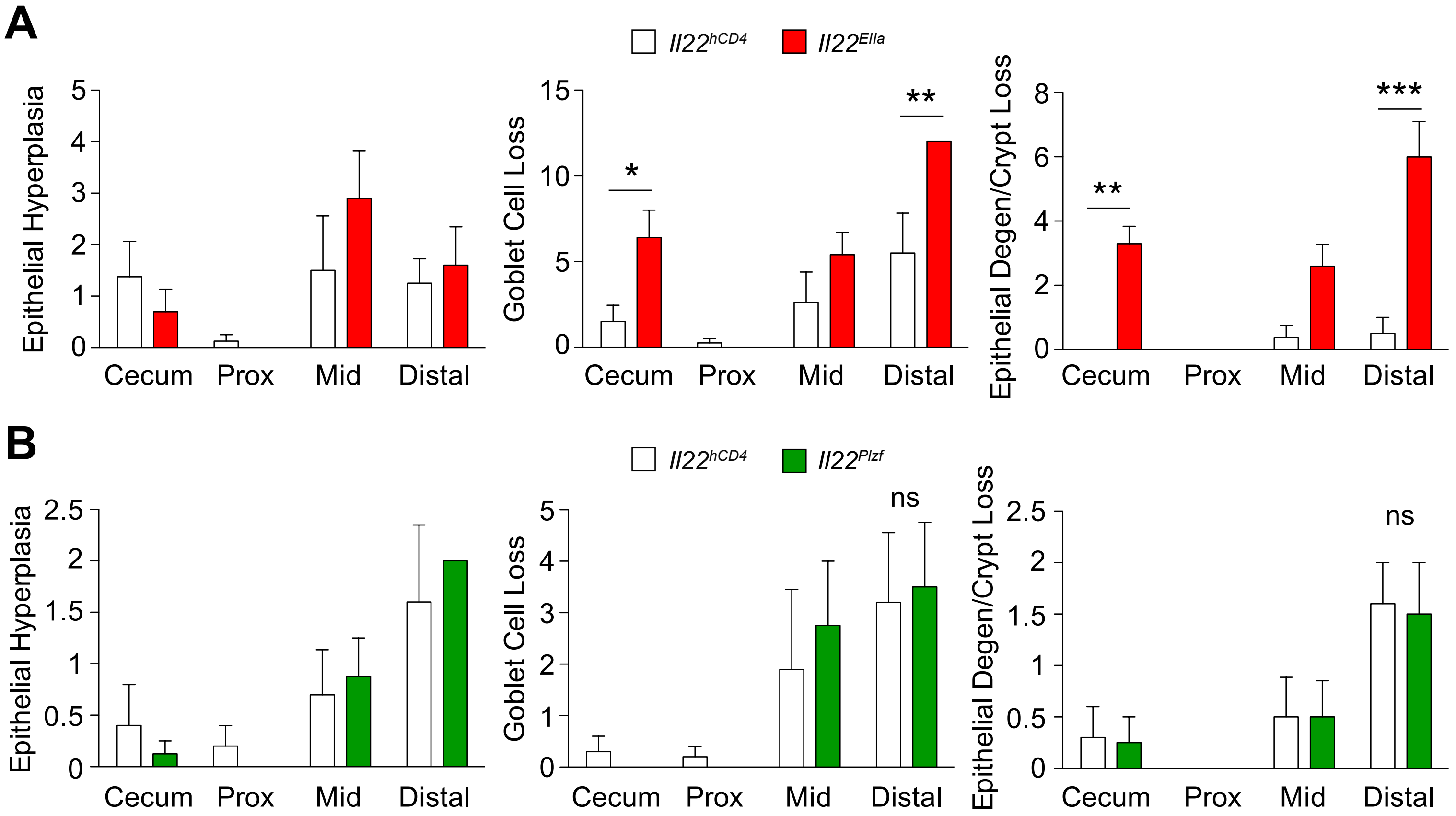

A
Il22hCD4
Il22EIIa
Epithelial Hyperplasia
Goblet Cell Loss
Epithelial Degen/Crypt Loss
Cecum
Prox
Mid
Distal
*
**
***
B
Il22hCD4
Il22Plzf
ns


**Figure S3**

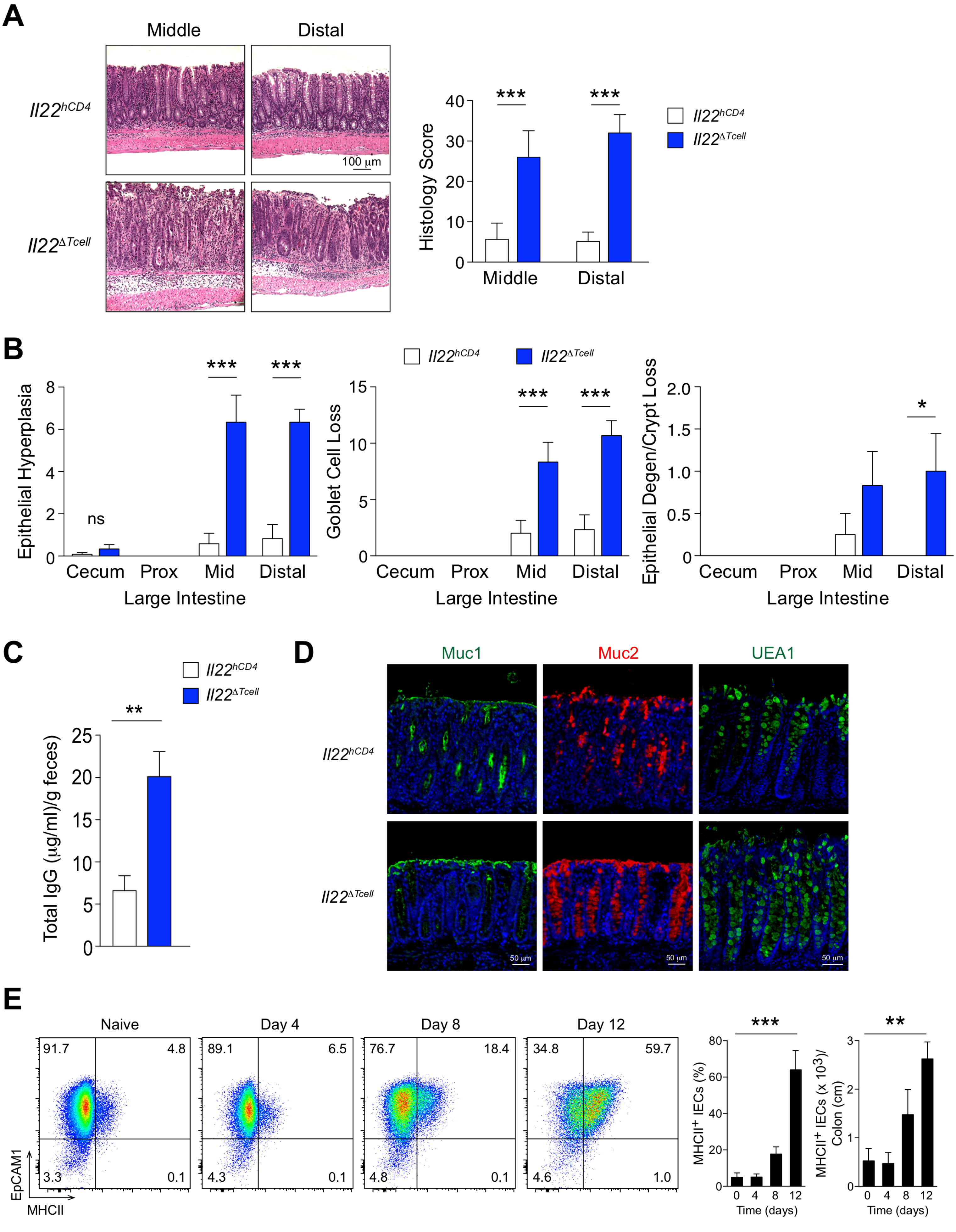
A
Middle
Distal
Il22hCD4
Il22ΔTcell
100 μm
Histology Score
40
30
20
10
0
***
***
Middle
Distal
Il22hCD4
Il22ΔTcell
B
Il22hCD4
Il22ΔTcell
Epithelial Hyperplasia
8
6
4
2
0
ns
***
***
Cecum
Prox
Mid
Distal
Large Intestine
Goblet Cell Loss
15
10
5
0
***
***
Cecum
Prox
Mid
Distal
Large Intestine
Epithelial Degen/Crypt Loss
2.0
1.5
1.0
0.5
0
*
Cecum
Prox
Mid
Distal
Large Intestine
C
Il22hCD4
Il22ΔTcell
**
Total IgG (μg/ml)/g feces)
25
20
15
10
5
0
D
Muc1
Muc2
UEA1
Il22hCD4
Il22ΔTcell
50 μm
50 μm
50 μm
E
Naive
Day 4
Day 8
Day 12
91.7
4.8
3.3
0.1
89.1
6.5
4.3
0.1
76.7
18.4
4.8
0.1
34.8
59.7
4.6
1.0
EpCAM1
MHCII
***
MHCII+ IECs (%)
80
60
40
20
0
0
4
8
12
Time (days)
**
MHCII+ IECs (x 10^3)/ Colon (cm)
3
2
1
0
0
4
8
12
Time (days)

**Figure S4**

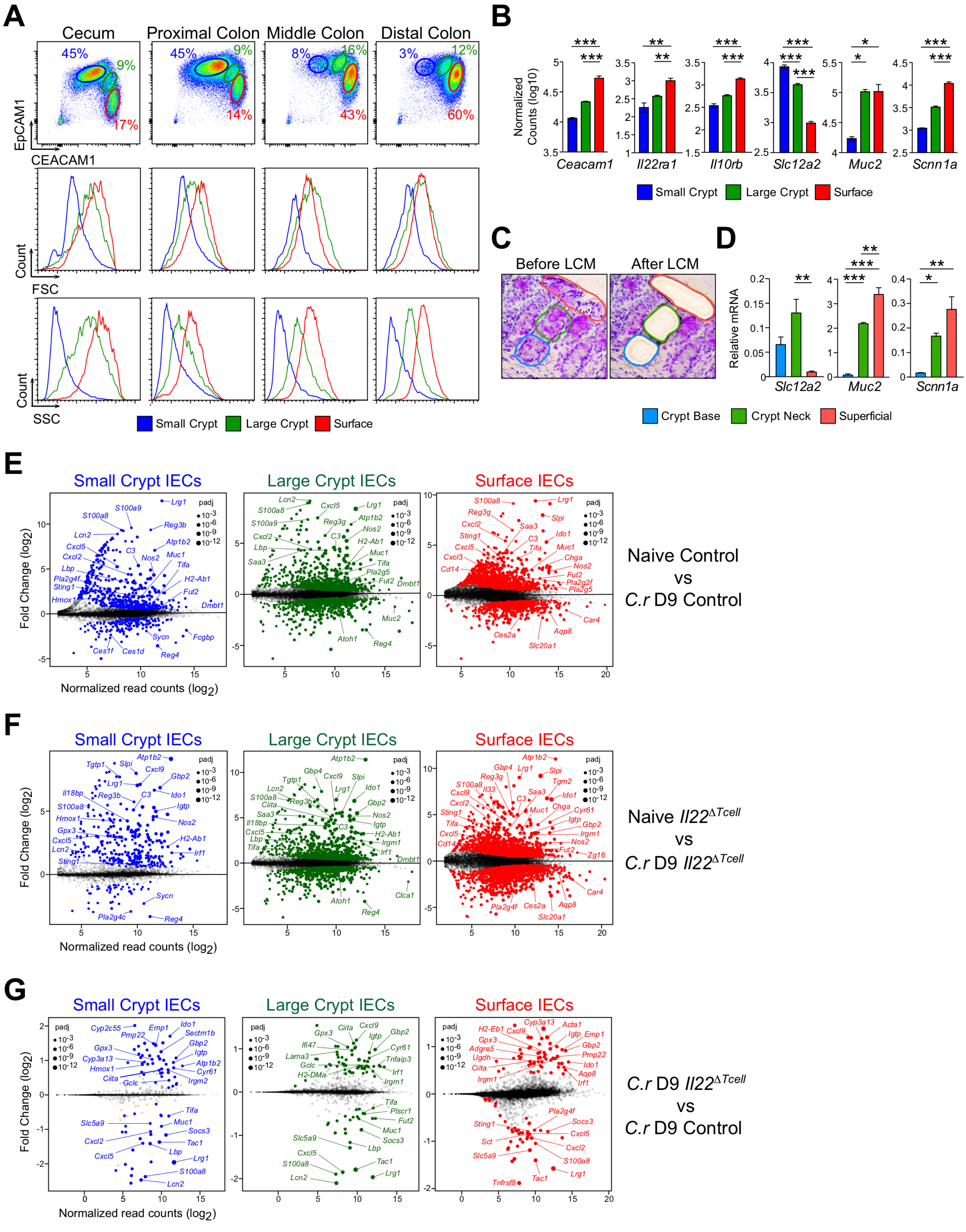

A
Cecum
Proximal Colon
Middle Colon
Distal Colon
EpCAM1
CEACAM1
Count
FSC
SSC
Small Crypt
Large Crypt
Surface
B
Normalized Counts (log10)
Ceacam1
Il22ra1
Il10rb
Slc12a2
Muc2
Scnn1a
C
Before LCM
After LCM
D
Relative mRNA
Crypt Base
Crypt Neck
Superficial
E
Small Crypt IECs
Large Crypt IECs
Surface IECs
Fold Change (log2)
Normalized read counts (log2)
padj
Naive Control
vs
C.r D9 Control
F
Naive Il22ΔTcell
vs
C.r D9 Il22ΔTcell
G
C.r D9 Il22ΔTcell
vs
C.r D9 Control


Figure S5

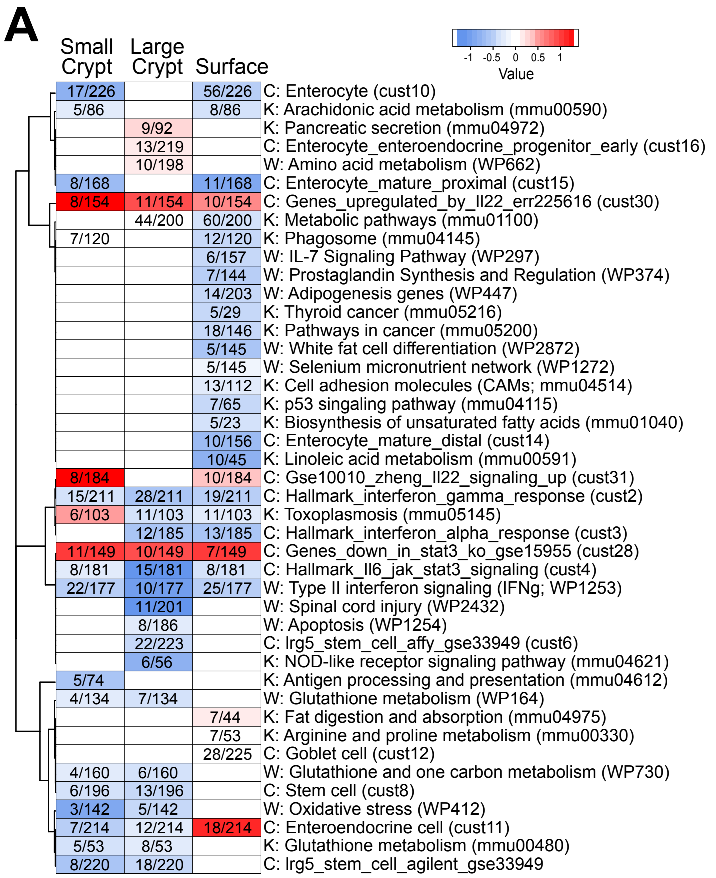


| Small Crypt | Large Crypt | Surface | |
|---|---|---|---|
| 17/226 | | 56/226 | C: Enterocyte (cust10) |
| 5/86 | | 8/86 | K: Arachidonic acid metabolism (mmu00590) |
| | 9/92 | | K: Pancreatic secretion (mmu04972) |
| | 13/219 | | C: Enterocyte_enteroendocrine_progenitor_early (cust16) |
| | 10/198 | | W: Amino acid metabolism (WP662) |
| 8/168 | | 11/168 | C: Enterocyte_mature_proximal (cust15) |
| 8/154 | 11/154 | 10/154 | C: Genes_upregulated_by_Il22_err225616 (cust30) |
| | 44/200 | 60/200 | K: Metabolic pathways (mmu01100) |
| 7/120 | | 12/120 | K: Phagosome (mmu04145) |
| | | 6/157 | W: IL-7 Signaling Pathway (WP297) |
| | | 7/144 | W: Prostaglandin Synthesis and Regulation (WP374) |
| | | 14/203 | W: Adipogenesis genes (WP447) |
| | | 5/29 | K: Thyroid cancer (mmu05216) |
| | | 18/146 | K: Pathways in cancer (mmu05200) |
| | | 5/145 | W: White fat cell differentiation (WP2872) |
| | | 5/145 | W: Selenium micronutrient network (WP1272) |
| | | 13/112 | K: Cell adhesion molecules (CAMs; mmu04514) |
| | | 7/65 | K: p53 singaling pathway (mmu04115) |
| | | 5/23 | K: Biosynthesis of unsaturated fatty acids (mmu01040) |
| | | 10/156 | C: Enterocyte_mature_distal (cust14) |
| | | 10/45 | K: Linoleic acid metabolism (mmu00591) |
| 8/184 | | 10/184 | C: Gse10010_zheng_Il22_signaling_up (cust31) |
| 15/211 | 28/211 | 19/211 | C: Hallmark_interferon_gamma_response (cust2) |
| 6/103 | 11/103 | 11/103 | K: Toxoplasmosis (mmu05145) |
| | 12/185 | 13/185 | C: Hallmark_interferon_alpha_response (cust3) |
| 11/149 | 10/149 | 7/149 | C: Genes_down_in_stat3_ko_gse15955 (cust28) |
| 8/181 | 15/181 | 8/181 | C: Hallmark_Il6_jak_stat3_signaling (cust4) |
| 22/177 | 10/177 | 25/177 | W: Type II interferon signaling (IFNg; WP1253) |
| | 11/201 | | W: Spinal cord injury (WP2432) |
| | 8/186 | | W: Apoptosis (WP1254) |
| | 22/223 | | C: Irg5_stem_cell_affy_gse33949 (cust6) |
| | 6/56 | | K: NOD-like receptor signaling pathway (mmu04621) |
| 5/74 | | | K: Antigen processing and presentation (mmu04612) |
| 4/134 | 7/134 | | W: Glutathione metabolism (WP164) |
| | | 7/44 | K: Fat digestion and absorption (mmu04975) |
| | | 7/53 | K: Arginine and proline metabolism (mmu00330) |
| | | 28/225 | C: Goblet cell (cust12) |
| 4/160 | 6/160 | | W: Glutathione and one carbon metabolism (WP730) |
| 6/196 | 13/196 | | C: Stem cell (cust8) |
| 3/142 | 5/142 | | W: Oxidative stress (WP412) |
| 7/214 | 12/214 | 18/214 | C: Enteroendocrine cell (cust11) |
| 5/53 | 8/53 | | K: Glutathione metabolism (mmu00480) |
| 8/220 | 18/220 | | C: Irg5_stem_cell_agilent_gse33949 |

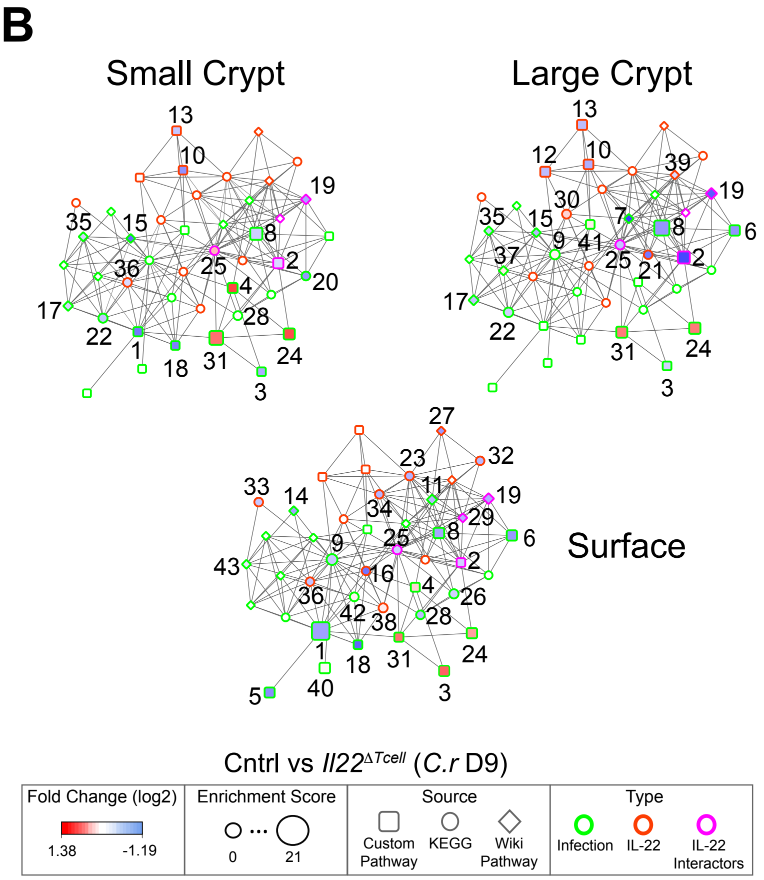


## C

Cntrl vs *Il22*$^{\Delta Tcell}$ (*C.r* D9)

| Ref_ID | Pathway | GS_ID | Pathway Name | Type | Fold Change (log2) Sm Crypt | Fold Change (log2) Lg Crypt | Fold Change (log2) Surface | Enrichment Score Sm Crypt | Enrichment Score Lg Crypt | Enrichment Score Surface | Final Score |
|---|---|---|---|---|---|---|---|---|---|---|---|
| 1 | Custom | cust10 | C: Enterocyte | Infection | -0.9 | 0.0 | -0.6 | 2.7 | 0.0 | 20.8 | 29.9 |
| 2 | Custom | cust4 | C: Hallmark_il6_jak_stat3_signaling | IL-22-interactor | -0.2 | -1.2 | -0.3 | 5.0 | 8.9 | 1.5 | 11.4 |
| 3 | Custom | cust11 | C: Enteroendocrine cell | Infection | -0.5 | -0.3 | 1.2 | 1.5 | 1.8 | 4.7 | 9.4 |
| 4 | Custom | cust31 | C: Gse10010_zheng_il22_signaling_up | Infection | 1.4 | 0.0 | 0.3 | 4.3 | 0.0 | 2.3 | 9.3 |
| 5 | Custom | cust14 | C: Enterocyte_mature_distal | Infection | 0.0 | 0.0 | -0.7 | 0.0 | 0.0 | 4.7 | 6.8 |
| 6 | Custom | cust3 | C: Hallmark_interferon_alpha_response | Infection | 0.0 | -0.7 | -0.6 | 0.0 | 5.1 | 5.1 | 6.8 |
| 7 | Wiki | WP2432 | W: Spinal Cord Injury | Infection | 0.0 | -1.2 | 0.0 | 0.0 | 2.3 | 0.0 | 5.5 |
| 8 | Custom | cust2 | C: Hallmark_interferon_gamma_response | Infection | -0.3 | -0.7 | -0.6 | 9.1 | 15.4 | 5.7 | 5.5 |
| 9 | KEGG | mmu01100 | K: Metabolic pathways | Infection | 0.0 | 0.1 | -0.3 | 0.0 | 3.9 | 6.3 | 4.4 |
| 10 | Custom | cust7 | C: Irg5_stem_cell_agilent_gse33949 | IL-22 | -0.7 | -0.5 | 0.0 | 1.4 | 3.8 | 0.0 | 4.1 |
| 11 | Wiki | WP447 | W: Adipogenesis genes | Infection | 0.0 | 0.0 | -0.4 | 0.0 | 0.0 | 4.4 | 3.9 |
| 12 | Custom | cust6 | C: Irg5_stem_cell_affy_gse33949 | IL-22 | 0.0 | -0.4 | 0.0 | 0.0 | 4.5 | 0.0 | 3.7 |
| 13 | Custom | cust8 | C: Stem cell | IL-22 | -0.4 | -0.4 | 0.0 | 1.8 | 4.6 | 0.0 | 3.1 |
| 14 | Wiki | WP374 | W: Prostaglandin Synthesis and Regulation | Infection | 0.0 | 0.0 | -0.6 | 0.0 | 0.0 | 2.5 | 2.9 |
| 15 | Wiki | WP412 | W: Oxidative Stress | Infection | -0.9 | -0.5 | 0.0 | 1.3 | 2.0 | 0.0 | 2.8 |
| 16 | KEGG | mmu00591 | K: Linoleic acid metabolism | IL-22 | 0.0 | 0.0 | -0.9 | 0.0 | 0.0 | 1.5 | 2.7 |
| 17 | Wiki | WP164 | W: Glutathione metabolism | Infection | -0.3 | -0.4 | 0.0 | 3.3 | 4.3 | 0.0 | 2.6 |
| 18 | Custom | cust15 | C: Enterocyte_mature_proximal | Infection | -0.8 | 0.0 | -1.0 | 1.3 | 0.0 | 1.6 | 2.6 |
| 19 | Wiki | WP1253 | W: Type II interferon signaling (IFNG) | IL-22-interactor | -0.5 | -1.0 | -0.5 | 3.7 | 6.5 | 5.7 | 2.6 |
| 20 | KEGG | mmu04612 | K: Antigen processing and presentation | Infection | -0.7 | 0.0 | 0.0 | 1.9 | 0.0 | 0.0 | 2.5 |
| 21 | KEGG | mmu04621 | K: NOD-like receptor signaling pathway | IL-22 | 0.0 | -0.8 | 0.0 | 0.0 | 1.5 | 0.0 | 2.5 |
| 22 | KEGG | mmu00480 | K: Glutathione metabolism | Infection | -0.4 | -0.3 | 0.0 | 2.7 | 3.5 | 0.0 | 2.2 |
| 23 | KEGG | mmu05200 | K: Pathways in cancer | IL-22 | 0.0 | 0.0 | -0.5 | 0.0 | 0.0 | 2.1 | 2.0 |
| 24 | Custom | cust30 | C: Genes_upregulated_by_il22_err225616 | Infection | 1.3 | 1.0 | 0.7 | 6.9 | 7.3 | 5.3 | 1.9 |
| 25 | KEGG | mmu05145 | K: Toxoplasmosis | IL-22-interactor | 0.5 | -0.4 | -0.3 | 1.9 | 3.1 | 2.3 | 1.8 |
| 26 | KEGG | mmu04514 | K: Cell adhesion molecules (CAMs) | Infection | 0.0 | 0.0 | -0.3 | 0.0 | 0.0 | 3.0 | 1.7 |
| 27 | Wiki | WP2872 | W: White fat cell differentiation | IL-22 | 0.0 | 0.0 | -0.6 | 0.0 | 0.0 | 1.4 | 1.7 |
| 28 | KEGG | mmu04145 | K: Phagasome | Infection | 0.0 | 0.0 | -0.4 | 1.9 | 0.0 | 1.8 | 1.6 |
| 29 | Wiki | WP297 | W: IL-7 Signaling Pathway | IL-22-interactor | 0.0 | 0.0 | -0.5 | 0.0 | 0.0 | 1.6 | 1.6 |
| 30 | KEGG | mmu04972 | K: Pancreatic secretion | IL-22 | 0.0 | 0.3 | 0.0 | 0.0 | 2.3 | 0.0 | 1.3 |
| 31 | Custom | cust28 | C: Genes_down_in_stat3_ko_gse15955 | Infection | 1.0 | 1.0 | 1.0 | 12.4 | 6.8 | 2.7 | 1.3 |
| 32 | KEGG | mmu05216 | K: Thyroid cancer | IL-22 | 0.0 | 0.0 | -0.4 | 0.0 | 0.0 | 1.5 | 1.3 |
| 33 | KEGG | mmu01040 | K: Biosynthesis of unsaturated fatty acids | IL-22 | 0.0 | 0.0 | -0.4 | 0.0 | 0.0 | 1.9 | 1.3 |
| 34 | KEGG | mmu04115 | K: p53 signaling pathway | IL-22 | 0.0 | 0.0 | -0.5 | 0.0 | 0.0 | 1.4 | 1.3 |
| 35 | Wiki | WP730 | W: Glutathione and one carbon metabolism | Infection | -0.3 | -0.3 | 0.0 | 1.9 | 2.1 | 0.0 | 1.2 |
| 36 | KEGG | mmu00590 | K: Arachidonic acid metabolism | IL-22 | -0.2 | 0.0 | -0.4 | 1.7 | 0.0 | 1.4 | 1.0 |
| 37 | Wiki | WP662 | W: Amino Acid metabolism | Infection | 0.0 | 0.2 | 0.0 | 0.0 | 2.6 | 0.0 | 0.9 |
| 38 | KEGG | mmu04975 | K: Fat digestion and absorption | IL-22 | 0.0 | 0.0 | 0.1 | 0.0 | 0.0 | 2.5 | 0.7 |
| 39 | Wiki | WP1254 | W: Apoptosis | IL-22 | 0.0 | -0.2 | 0.0 | 0.0 | 1.5 | 0.0 | 0.7 |
| 40 | Custom | cust12 | C: Goblet cell | Infection | 0.0 | 0.0 | 0.0 | 0.0 | 0.0 | 5.7 | 0.5 |
| 41 | Custom | cust16 | C: Enterocyte_enteroendocrine_progenitor_early | Infection | 0.0 | 0.1 | 0.0 | 0.0 | 1.6 | 0.0 | 0.5 |
| 42 | KEGG | mmu00330 | K: Arginine and proline metabolism | Infection | 0.0 | 0.0 | 0.1 | 0.0 | 0.0 | 2.0 | 0.4 |
| 43 | Wiki | WP1272 | W: Selenium Micronutrient Network | Infection | 0.0 | 0.0 | -0.1 | 0.0 | 0.0 | 1.4 | 0.4 |

**Figure S6**
